# Peripersonal encoding of forelimb proprioception in the mouse somatosensory cortex

**DOI:** 10.1101/2022.05.25.493376

**Authors:** Ignacio Alonso, Irina Scheer, Mélanie Palacio-Manzano, Noémie Frézel-Jacob, Antoine Philippides, Mario Prsa

**Author notes:** equal contribution (Ignacio Alonso, Irina Scheer).

## Abstract

Conscious perception of limb movements depends on proprioceptive neural responses in the somatosensory cortex. In contrast to tactile sensations, proprioceptive cortical coding is barely studied in the mammalian brain and practically non-existent in rodent research. To understand the cortical representation of this important sensory modality we developed a passive forelimb displacement paradigm in behaving mice and also trained them to perceptually discriminate where their limb is moved in space. We delineated, for the first time, the rodent proprioceptive cortex with wide-field calcium imaging and optogenetic silencing experiments during awake behavior. Our results reveal that proprioception is represented in both sensory and motor cortical areas. In addition, behavioral measurements and responses of layer 2/3 neurons imaged with two-photon microscopy reveal that passive limb movements are both perceived and encoded in the mouse cortex as a spatial direction vector that interfaces the limb with the body’s peripersonal space.

## Introduction

Proprioception allows us to detect and track the movements of our limbs. The major target of proprioceptive signals is the cerebellum (spino- and cuneo-cerebellar tracts), involved in maintaining limb posture and adapting movements to unexpected perturbations; processes that typically occur subconsciously ^1^. Sensory afferents also ascend to the cerebral cortex (dorsal column-medial lemniscus pathway) where proprioceptive information is consciously perceived^1,2^. How the proprioceptive sensation of a limb movement is perceived and encoded by neurons in the somatosensory cortex (S1) is still poorly understood. Practically all functional studies of S1 in rodents use extracorporeal (i.e. tactile) stimuli and the few studies in primates on limb proprioception examined cortical responses mainly to active reaching movements^3–6^. The sensation of an active limb movement is however strongly dominated by motor signals^7,8^. During muscle contractions, gamma motor neurons tune the sensitivity of both muscle and joint proprioceptors in a manner that is still not fully understood^9,10^. Cortical sensory responses are in addition modulated by motor efference copies during active movements^5^, which therefore reveal little about how neurons in S1 encode proprioceptive ex-afference on its own. Studying limb movements in the absence of muscle contraction and predictive processing^11^ is needed to understand the contribution of ascending sensory signals to the cortical proprioceptive code.

In this study, we conducted a series of anatomical tracing, behavioral, neuronal imaging and optogenetic manipulation experiments to investigate where and how passive forelimb movements are represented by the activity of neurons in the mouse cortex and how these signals are perceived. Previous similar experiments in primates are few^3,5,6,12^, based on a limited range of stimuli, and do not assess their perceptual significance. We applied well-controlled and reliable proprioceptive stimuli using a robotic manipulandum and trained mice in a new perceptual discrimination task. This allowed us to locate the proprioceptive cortex and examine the relationship between spatial variables, perception and cortical neuronal activity. Our findings provide the first characterization of perceptually relevant neural encoding of proprioception.

## Results

### Conscious proprioceptive pathway in the mouse brain

The most direct route for proprioceptive inputs to reach the cerebral cortex is via the dorsal column nuclei^2,13^. Afferents from forelimb proprioceptors ascend in the dorsal column and synapse onto second order neurons in the external cuneate nucleus (ECu), which project primarily to the cerebellum^13,14^ (i.e. the “non-conscious” pathway). In addition to the cerebellar projection, a thalamic projection from the ECu has been confirmed in primates^15^, raccoons^16^ and rats^17^, but seems not be present in cats^18^. It has thus been suggested that this “conscious” pathway is phylogenetically recent and important for dexterous limb movements^13^.

To determine whether a direct proprioceptive route from the forelimb to the cerebral cortex also exists in mice, we first generated PV-Cre; Ai32 mice, in which parvalbumin (PV), a reliable neurochemical marker for proprioceptors^19,20^, is labeled with a green fluorescent protein. We then genetically restricted the expression of the red fluorescent protein tdTomato (tdTom) to proprioceptive forelimb afferents by means of AAV9-flex-tdTom injections in the biceps brachii and triceps brachii long muscles (Fig. 1A). This was confirmed by detecting PV and tdTom co-labeling of their cell bodies in cervical dorsal root ganglia (Fig. 1B). By observing the innervation patterns of their central branches in immunostained sections of the medulla (Fig. 1C), we show that primary proprioceptive neurons of the forelimb mainly innervate the ECu, and thus confirm their segregation from tactile afferents that terminate in the adjacent cuneate nucleus (Cu)^20,21^. To test whether axons from ECu neurons ascend towards the cortex, we next retrogradely labeled cuneo-thalamic projections. We targeted injections of AAVretro-tdTom and AAVretro-eGFP to ventral posterior lateral (VPL) and posterior (PO) thalamic nuclei (Fig. 1D), which relay lemniscal afferents to the forelimb somatosensory cortex (fS1)^22^. In addition to the expected labeling in Cu, we observed large labeled cells characteristic of ECu neurons^13^ located dorsolaterally to the Cu (Fig. 1E). Therefore, second order neurons in the dorsal column nuclei of mice also seem to convey forelimb proprioceptive signals along the so-called conscious proprioceptive pathway.

**Figure 1.**
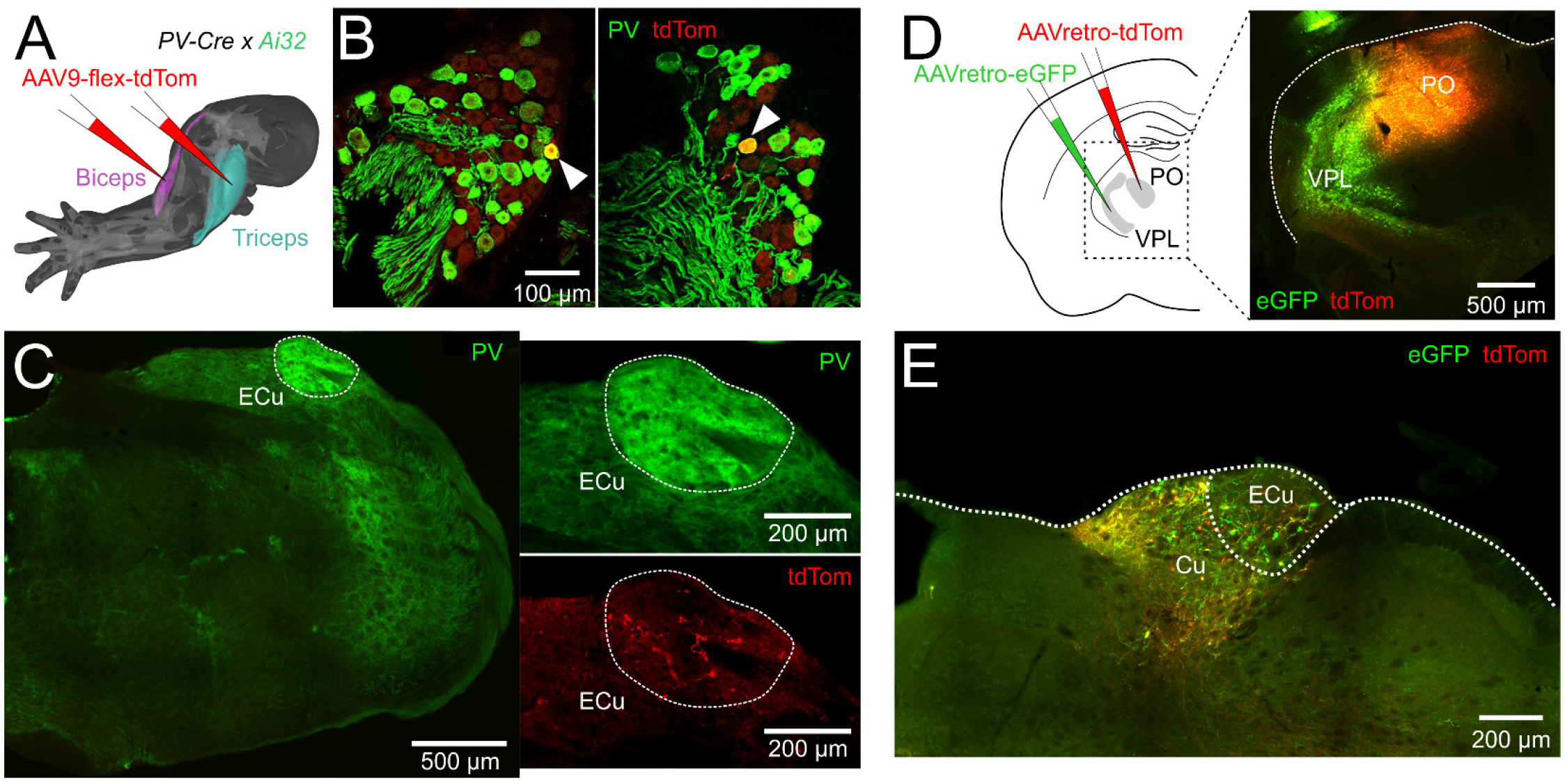
Forelimb proprioceptive afferents ascend to the mouse cortex via the cuneo-thalamic pathway. **A:** Genetically restricted labeling of proprioceptive afferents from forelimb muscles with AAV9-flex-tdTomato in PV-Cre;Ai32 mice. **B:** Confirmation of labeled PV+ cell bodies of primary sensory neurons in the cervical DRG (arrows). **C:** Central branches of the labeled pseudo-unipolar DRG neurons terminate in the ECu. **D:** Retrograde labeling of cuneo-thalamic projections with AAVretro-tdTomato and AAVretro-GFP injections in PO and VPL of the sensory thalamus. **E:** Labeled thalamus projecting neurons are found in both Cu and ECu.

### Perceptually relevant proprioceptive signals activate both sensory and motor cortical areas

In primates, the cortical recipient of proprioceptive afferents from muscles (also tendons and joints) is Brodmann’s area 3a, architectonically distinct from the adjacent somatosensory area 3b receiving cutaneous inputs^2,23^. The mouse fS1 has been previously studied in terms of its tactile responses^24,25^ and is thus typically thought of as homologous to area 3b. Cutaneous and proprioceptive afference from the forelimb innervate separate dorsal column nuclei; Cu and ECu, respectively^20,21^. The two pathways overlap but continue to be distinct in thalamic nuclei^17,26^. It follows that, like in primates, limb proprioception might be represented separately from touch in the mouse cortex and therefore not limited to fS1. Whether the rodent somatosensory cortex has a proprioceptive area (i.e. a homologue of area 3a) distinct from fS1 is unknown^27^. To address this question, we generated Rasgrf2-dCre; Ai148 mice expressing the Ca^2+^ indicator GCaMP6f in cortical layer 2/3 neurons and trained them in a new proprioceptive stimulation task.

Recent publications highlight the primacy of studying neural circuits during well-quantifiable behavioral tasks^28,29^ instead of anesthetized animals as is often the case with sensory mapping studies. Such tasks are already well established for practically all sensory modalities but proprioception. To our knowledge, no paradigm currently exists in rodents for stimulation of proprioception in the awake behaving condition. We have therefore developed a method for systematic and quantifiable delivery of proprioceptive stimuli to the mouse forelimb, which is amenable to simultaneous imaging of cortical activity (i.e. under head fixation). Mice were trained to grasp the endpoint of a robotic manipulandum (Fig. 2A, Supplementary Fig. 1A) and allow their right forelimb to be passively displaced in any of 8 co-planar horizontal directions. In each trial, the passive movement displaced the limb from the home to the target position and, following a random time delay, back to home (Fig. 2B, Supplementary Movie 1). Continuous holding throughout all epochs resulted in a correct trial and a water droplet reward. Otherwise, the trial was aborted and the mouse punished (air puff) immediately upon releasing the manipulandum. We simultaneously imaged large-scale Ca^2+^ dependent neocortical activity with a wide-field fluorescence macroscope through a transparent skull preparation (Fig. 2A). The activation pattern evoked by the proprioceptive stimulation of the forelimb (correct trials) covered the entire anterolateral extent of contralateral fS1 but also extended medially to the primary motor cortex (M1) (Fig. 2C). As a control, vibrotactile stimulation of the paw (100 Hz vibration applied to the endpoint holder) did not activate motor areas in a similar way (Fig. 2C,D). Also, the location of peak activity was highly consistent across mice (N=7) for tactile but not for proprioceptive stimulation (Fig. 2D).

**Figure 2.**
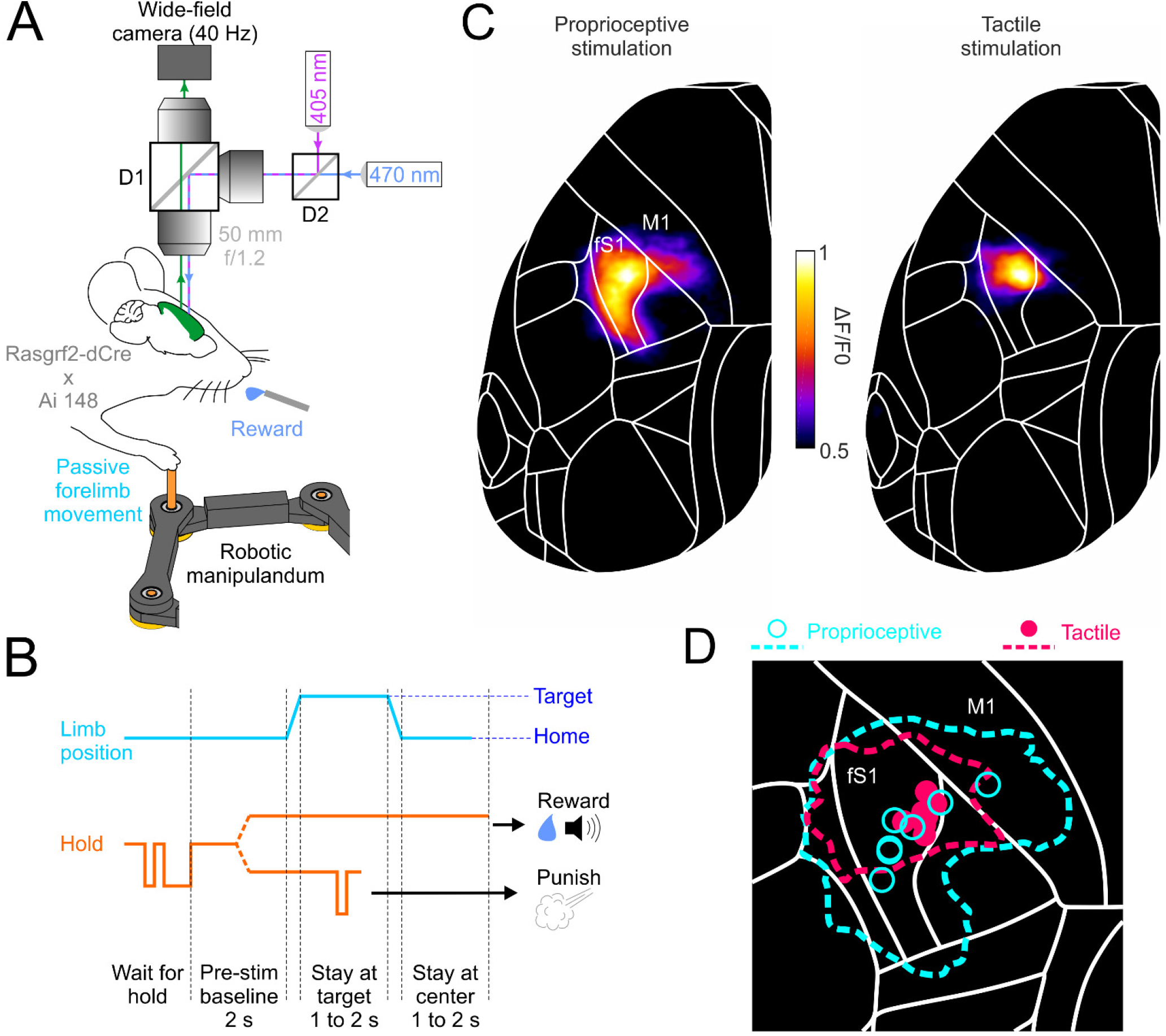
Cortex-wide imaging of neural activity during proprioceptive stimulation of the mouse forelimb. **A:** Schematic of the macroscope (D1, D2: dichroic mirrors) for wide-field imaging of Ca2+ dependent cortical activity in Rasgrf2-dCre; Ai148 mice during passive forelimb displacements with a robotic manipulandum. **B:** Trial timeline of the passive forelimb displacement task (split dotted lines denote different trial outcomes). **C:** Mean cortical activation maps (n=7 mice) in the contralateral hemisphere (registered to the 2D top projection of the Allen Mouse Brain Atlas; mouse.brain-map.org) produced by proprioceptive and tactile stimuli. **D:** Mean activation map contours (within 50% of peak activity) and peak activity loci (symbols) of individual mice (n=7).

The mouse proprioceptive cortex thus does not seem to be limited to fS1, but also encompasses the medially adjacent M1, known as the caudal forelimb motor area (CFA)^30^. Are proprioceptive signals in both fS1 and CFA necessary for conscious perception of limb movements? Might either of them instead underlie proprioceptive processes that occur subconsciously (e.g. reflex mediation, feedback control of movement, limb coordination etc.)? To assess perceptual relevance, we trained VGAT-ChR2 mice to discriminate between two proprioceptive stimuli (lateral vs. medial forelimb displacement). Because these mice express the light-gated ion channel ChR2 in GABAergic neocortical neurons, we could optogenetically silence small areas of cortex during behavior through a transparent skull preparation (Fig. 3A). The mice performed a two-alternative forced choice task (lick left vs. right, Supplementary Movie 2), which included a delay period to temporally separate and eliminate confounds between sensory (stimulus epoch) and motor (response epoch)x processes during inactivation^31^ (Fig. 3B, see Methods for details). Expert mice (N=4) performed the discrimination task at a high level of accuracy (>75% correct for displacements >= 3mm, Fig. 3C); significant discrimination was observed for displacements as small as 1 mm (p<0.05, binomial test, two-sided). Silencing of contralateral fS1, but not control sensory areas (hindlimb, whisker or ipsilateral forelimb S1), significantly decreased the percentage of correct answers compared to baseline trials without cortical inactivation (Fig. 3D). In accordance with the observed GCaMP activation patterns, silencing of CFA also significantly decreased performance. The same was not observed when we targeted the anterior lateral motor cortex (ALM), a pre-motor like area involved in preparatory motor activity^32^. We therefore exclude the possibility that CFA inactivation affected motor instead of sensory/perceptual aspects of the task. In fact, the effects of ALM silencing on behavior became apparent with shorter durations of the delay period (<= 500 ms, Fig. 3E). This result indicates that our delay period of 1 s effectively postponed the preparation of the motor response and dissociated it from the stimulus epoch.

**Figure 3.**
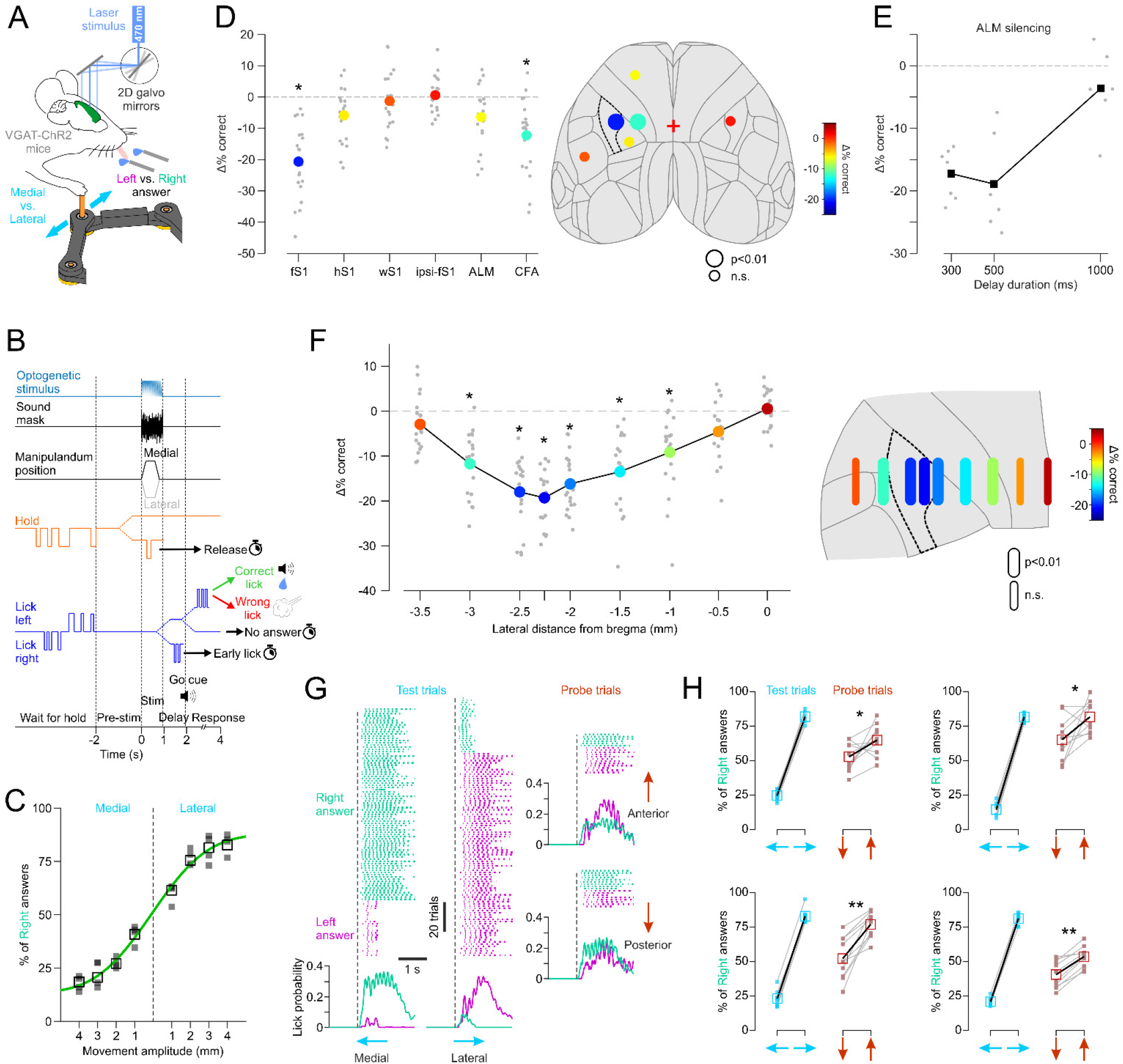
Identification of behavioral variables and cortical areas necessary for perceptual discrimination of forelimb proprioception. **A**: Schematic of the optogenetic silencing experiment during a 2AFC proprioceptive discrimination task. **B**: Trial timeline (split dotted lines denote different trial outcomes). **C**: Psychometric discrimination curve fitted to the mean (empty squares, minimum 200 trials/amplitude) answer % (N=4 mice, filled squares). **D:** Left, mean (colored circles, N=4 mice) decrease in performance (difference in % of correct trials compared to no stimulation) during selective optogenetic silencing (single point stimulation) of different cortical areas. Grey circles are data from individual sessions (5 sessions per mouse per area). fS1: forelimb S1, hS1: hindlimb S1, wS1: whisker S1, ipsi-fS1: ipsilateral fS1, ALM: anterior lateral motor area, CFA: caudal forelimb motor area. *: p<0.01 (two-sided t-test, Bonferroni corrected). Right, same data depicted the top projection of the Allen Mouse Brain Atlas (fS1 is highlighted by dotted lines). **E:** decrease in performance during ALM silencing with different delay durations (1 mouse; black squares: means; grey circles: individual sessions). **F:** Left, decrease in performance (same statistics as in D) during silencing (1 mm line stimulation) of contralateral cortical areas at the same anterio-posterior location (corresponding to fS1) but different lateral distances relative to bregma. Right, same data depicted on a zoomed in projection of the Allen Mouse Brain Atlas. **G:** Example session showing lick events on each trial (colored ticks sorted by right vs. left answer) and instantaneous lick probability of each spout (based on all trials) for trained (medial and lateral) and probe (anterior and posterior) stimuli. **H:** mean % of right answers (empty squares and black lines) for individual mice (different panels) of 10 sessions (small filled symbols and grey lines) when 15% of trials were probe stimuli. Anterior movements resulted in significantly more right answers than posterior movements in all 4 mice (*: p<0.05, **: p<0.01, two-sided t-test).

We next asked whether a perceptually relevant proprioceptive “hotspot” exists in the mouse sensorimotor cortex. Indeed, the transitional zone between fS1 and CFA, but also the dysgranular zone between fS1 and the more lateral orofacial somatosensory cortex, have been hypothesized to be the rodent homologue of area 3a^23,33–36^. If this “hotspot” exists, we expect its inactivation to have the strongest effect on discrimination ability. We therefore inactivated 1 mm strips of cortex centered 0.25 mm anterior and between 0 and −3.5 mm lateral to bregma in 0.5 mm increments. The strongest inactivation effects were observed between −2.5 and −2 mm (Fig. 3F), which, after registering the coordinates for each mouse to the Allen Mouse Brain Atlas (Supplementary Fig. 2), corresponds to the more medial end of fS1. We also observed that correct performance decreases more when silencing medially toward motor cortex than laterally towards orofacial somatosensory areas (Fig. 3F). We conclude that, rather than being a distinctly defined unit like the primate area 3a, the primary proprioceptive cortex in mice has a diffuse representation across S1 and M1.

### Limb proprioception is perceived in relation to the body

What do the mice actually perceive when they perform the discrimination task (i.e. to identify lateral vs. medial limb displacements)? To answer this question, in expert mice (10 sessions per mouse with >75% correct answers), we introduced probe stimuli randomly on 15% of trials. The probe stimuli were limb displacements in the anterior or posterior direction and were rewarded regardless of the answer (lick left vs. right). Because both are spatially equidistant from the two trained test stimuli (movements in the posterior and anterior directions are both 90 degrees apart from either the lateral or medial directions), a perceptual association between probe and test stimuli based on proximity in allocentric coordinates is not possible. We nevertheless observed that anterior and posterior probe trials did not result in the same ratio of left vs. right answers (Fig. 3G). Answers to anterior displacements were more similar to lateral stimuli and those to posterior displacements to medial stimuli. The difference in % of right vs. left answers was expectedly smaller than for test trials, but this perceptual bias was highly consistent and observed in all 4 mice (Fig. 3H).

Why did mice make this particular perceptual association between trained and neutral stimuli? One explanation is that the associated displacements produce more similar changes in joint angles. Indeed, lateral vs. medial forelimb displacement can also be described as an adduction vs. abduction of the humerus. We therefore need to quantify how the humeral angle with the earth vertical axis changes with movement in different directions. Joint tracking is problematic in the mouse forelimb given the absence of clear visual features. The proximal part of the limb is covered by a large volume of skin and subcutaneous adipose tissue which are loosely connected to bones and they do not move in conjunction as a result. The locations of shoulder, elbow and scapulothoracic joints are thus hidden and cannot be identified using standard video tracking methods. To overcome this problem, we made an ex-vivo surgical preparation allowing the identification of joint positions on images of the mouse musculature (see Methods for details). Images of the limb, as it was displaced by the manipulandum throughout the planar workspace (Supplementary Fig. 1B), were acquired by a stereo camera system and, after triangulation, allowed extraction of their 3D coordinates (Fig. 4A and Supplementary Movie 3, see Methods for details). These measurements are not a precise quantification of joint locations in vivo, nor do they account for any existing mouse to mouse variability in posture or limb impedance. The 3D reconstruction is however a good approximation of how joint angles generally change for movements in different directions. We specifically calculated the humerus adduction/abduction angle and mapped it onto the planar movement workspace (Fig. 4A). Relative changes of the joint angle resulting from any movement within the workspace could thus be read from the obtained map. The map shows that lateral and posterior stimuli result in an abduction of the humerus, whereas anterior and medial limb movements produce an adduction (Fig. 4B). This similarity in joint angles therefore cannot explain the perceptual association that we observed (Fig. 4C). Instead, we suggest that mice perceived whether the limb’s endpoint was being displaced away from the body (in lateral and anterior directions) or towards the body (in medial and posterior directions). Accordingly, limb proprioception might be primarily perceived in terms of body-fugal vs. body-petal movements.

**Figure 4.**
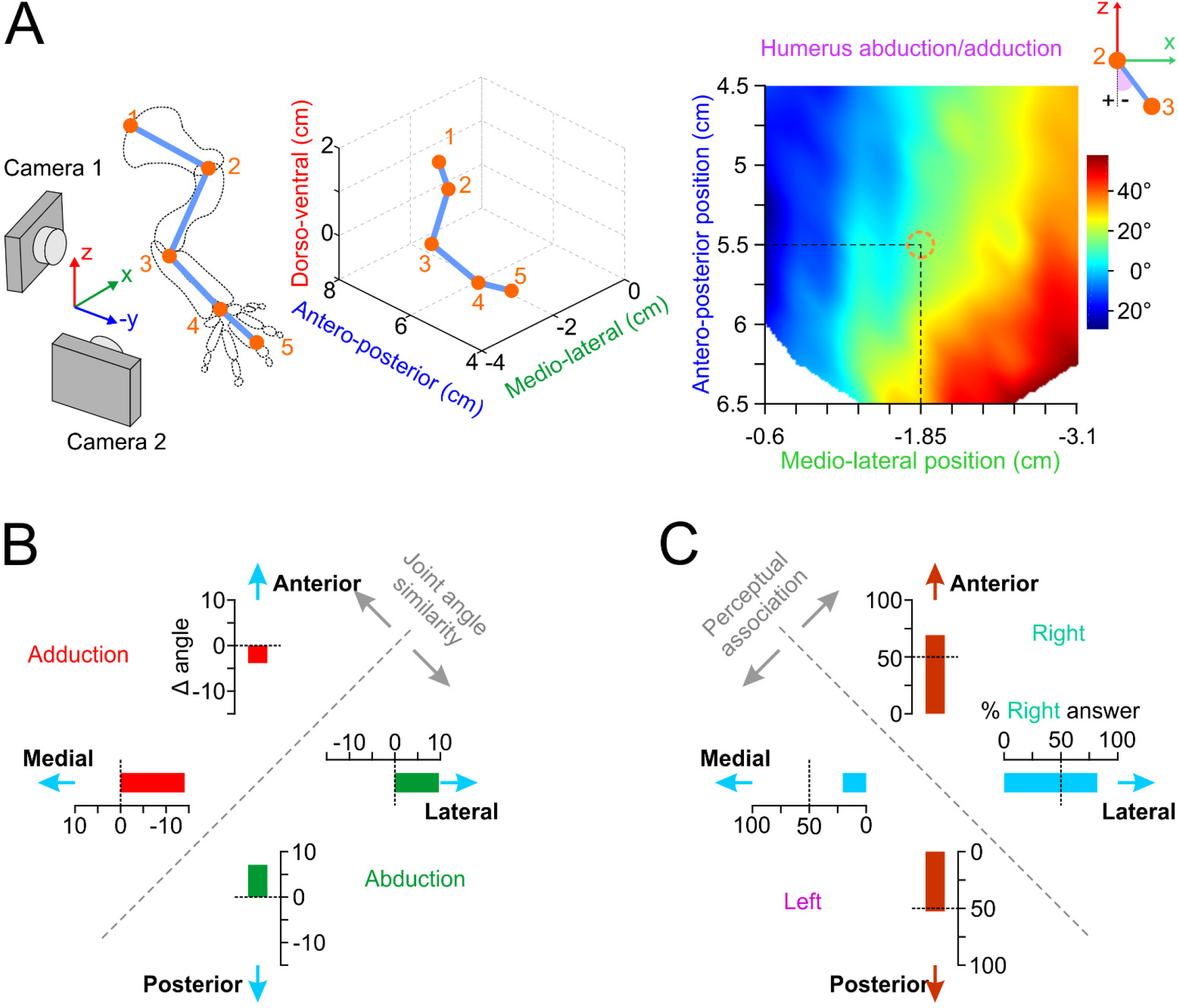
Perceptual discrimination is not based on changes in joint angles. **A:** Identification and tracking of mouse forelimb joints with stereo cameras (see Methods for details) yields the humerus abduction/adduction angle (color map, linear interpolation) mapped onto the planar workspace defined in Supplementary Fig. 1B (orange circle: manipulandum’s home position). **B:** Changes (Δ angle measured from the map in A) in humerus adduction (negative values) or abduction (positive values) as the limb is displaced by 4 mm in the medial, lateral, posterior and anterior directions. **C:** % right answer means (N=4 mice) of the data in Fig. 3H for the trained (medial and lateral) and probe (anterior and posterior) directions. The perceptual association axis is orthogonal to the joint similarity axis in B.

### Proprioceptive neurons in mouse forelimb somatosensory cortex

Is there a neural correlate of a body-fugal vs. body-petal representation of forelimb proprioception in the mouse cortex? To address this question, we imaged with two-photon microscopy the Ca^2+^ dependent activity of neurons in fS1 during the robotic forelimb displacement task (Fig. 5A). The imaged neurons most often responded to the stimuli in a phasic manner (i.e. to the dynamic component of the movement). Occasionally, we also observed sustained responses (tonic or both phasic and tonic) when the forelimb was being held at the target position (Fig. 5B). These three response types are also characteristic of how muscle spindle afferents respond to passive muscle stretch^37,38^. To assess possible contamination by tactile or motor signals, we compared how the neurons respond to active touch and active release events during the pre-stimulus period. Responses to release events were rarely greater, and responses to touch events never greater than those to passive movement (Fig. 5C, D). In addition, we tested how the neurons respond to passive tactile stimulation of the forepaw. There was virtually no overlap between neurons activated by forelimb displacement and those activated by tactile stimulation of the glabrous forepaw skin (Fig. 5E). Furthermore, pharmacologically blocking sensory afference from the paw had no significant effect on responses to limb movement, whereas it strongly suppressed tactile responses (Fig. 5F). We conclude that the neurons responsive to passive forelimb movements we imaged in fS1 are mainly driven by proprioceptive sensory inputs.

**Figure 5.**
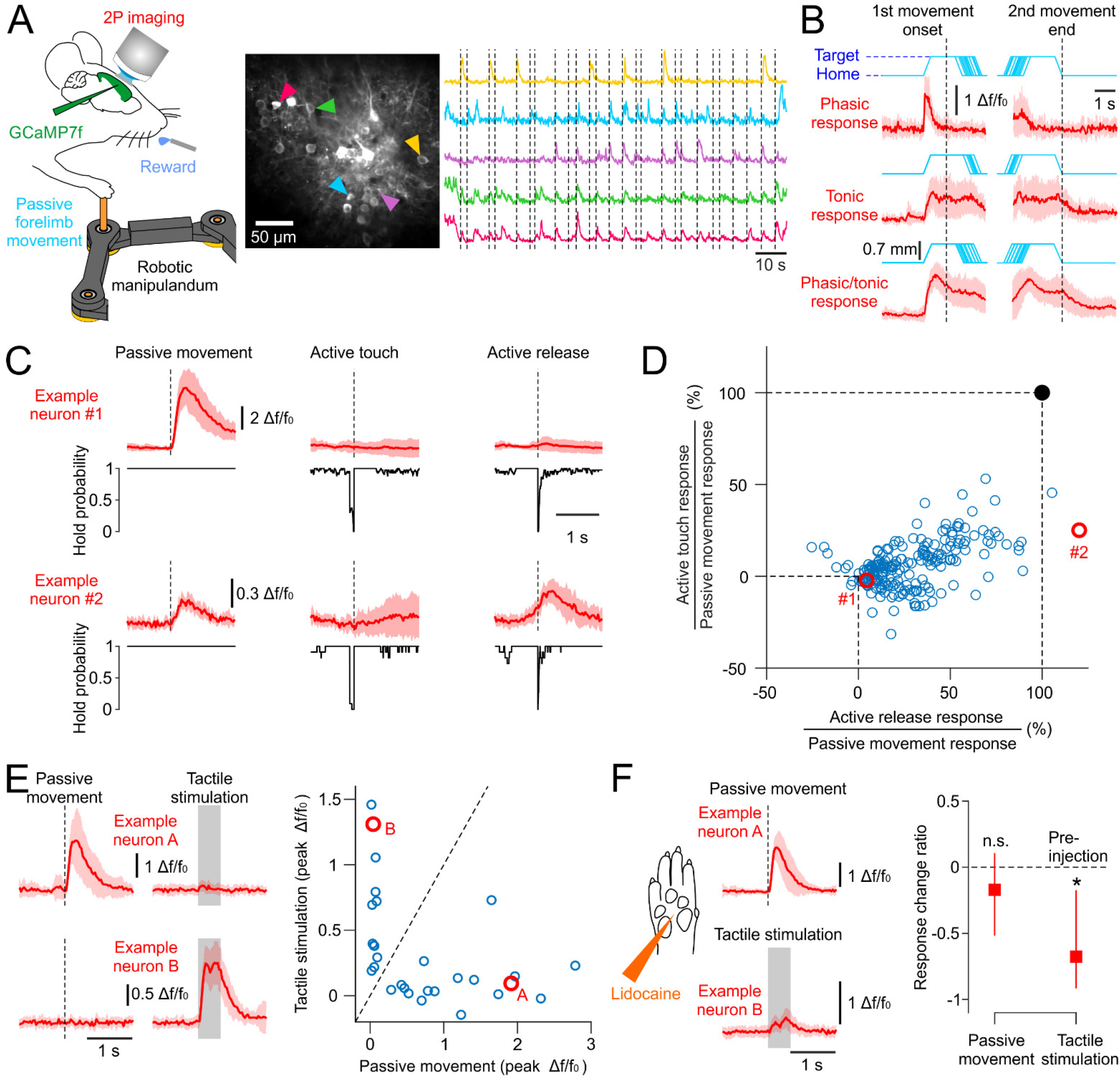
Ca^2+^ imaging of proprioceptive neuronal responses in the mouse forelimb S1. **A**: Left, experiment schematic of the passive forelimb displacement task with simultaneous two-photon imaging of cortical neurons transfected with GCaMP. Right, cropped two-photon image of the forelimb somatosensory cortex and Ca^2+^ dependent activity traces of 5 neurons responding at the onset of passive forelimb movements (dotted lines, 8 different directions tested). **B**: Three types of observed proprioceptive responses. *Δf/f_0_* mean (± s.d.) traces of three example neurons (red) aligned to first movement (home-to-target) onset and second movement (target-to-home) end (cyan: individual movement trajectories). **C**: Mean (± s.d.) responses (red) of two example neurons to passive forelimb movement, active touch and active release of the manipulandum. Black traces: instantaneous probability to hold the manipulandum across trials. **D**: Peak responses to active touch and active release as a % of peak responses to passive movement (N=238 neurons, 18 mice). Red circles: data of example neurons in C. **E**: Left, Mean (± s.d.) responses (red) of two example neurons to passive forelimb movement and tactile stimulation of the paw glabrous skin (shaded rectangle indicates the duration of skin indentation). Right, Peak responses of 29 neurons (2 mice) to passive movement vs. tactile stimulation. Red circles: data of example neurons in the left panel. **F**: Responses after nerve block (s.c. lidocaine injection in the paw) of the two example neurons and the response change ratio of the imaged population (median ± quartiles) relative to their pre-injection levels (N=14 neurons for passive movement, N=12 neurons for tactile stimulation, 2 mice). *: p<0.01 n.s.: p=0.58 (Wilcoxon signed rank test, two-sided).

### Directional selectivity of proprioceptive neurons reveals a peripersonal representation

Are proprioceptive neurons in fS1 directionally selective and can their selectivity account for our behavioral results (Fig. 3H, 4C)? In the classic studies in primates, the activity of neurons across motor^39,40^ and somatosensory^6^ cortical areas was found to be tuned to the spatial direction of active reaching movements. Similar to motor directional tuning, selectivity for directional sensory stimuli is characteristic of neurons in sensory cortices^41,42^. Data for directional selectivity of somatosensory cortex neurons to passive arm movements in primates is limited and often tested with poorly quantified stimuli (e.g. short arm perturbations or bumps)^3,5,6^, and is to our knowledge non-existent in rodents.

We imaged fS1 neurons in mice during passive displacements of their forelimbs in eight co-planar directions. The robotic manipulandum produced highly consistent trajectories and movement kinematics in the eight directions (Supplementary Fig. 1C, D), which were therefore unaffected by the impedance of the mouse limb. The activity of almost all responsive cells (>95%) was directionally selective (Fig. 6A, B); their activity could be expressed as a Gaussian function of movement direction (226/238 cells with significant fits). Interestingly, their preferred directions were not uniformly distributed (p<0.01, Rayleigh test) across the targeted space. The majority of neurons preferred movements to targets posterior and medial to the home position, and very few to movements in the anterior and lateral directions (Fig. 6C). The non-uniformity is unlikely due to a sampling bias (imaging locations biased to a particular area of fS1 where certain directions might be overrepresented) given that the imaged neurons covered the whole antero-posterior extent of fS1 and no obvious directional topography could be observed (Supplementary Fig. 3). The same directional preference was observed for home-to-target as for target-to-home movements (Fig. 6A,C, cyan and magenta arrows, respectively), that is for identical movement vectors that displace the limb through different spatial positions. Indeed, the angular shift between preferred spatial positions for home-to-target and target-to-home movements was normally distributed around 180° (Fig. 6D) meaning that directional preference was preserved. It follows that proprioceptive fS1 neurons are responsive to movement direction per-se rather than driven by the limb crossing a particular spatial location; they are not postural or place cell like representations of a body part in S1^43^.

**Figure 6.**
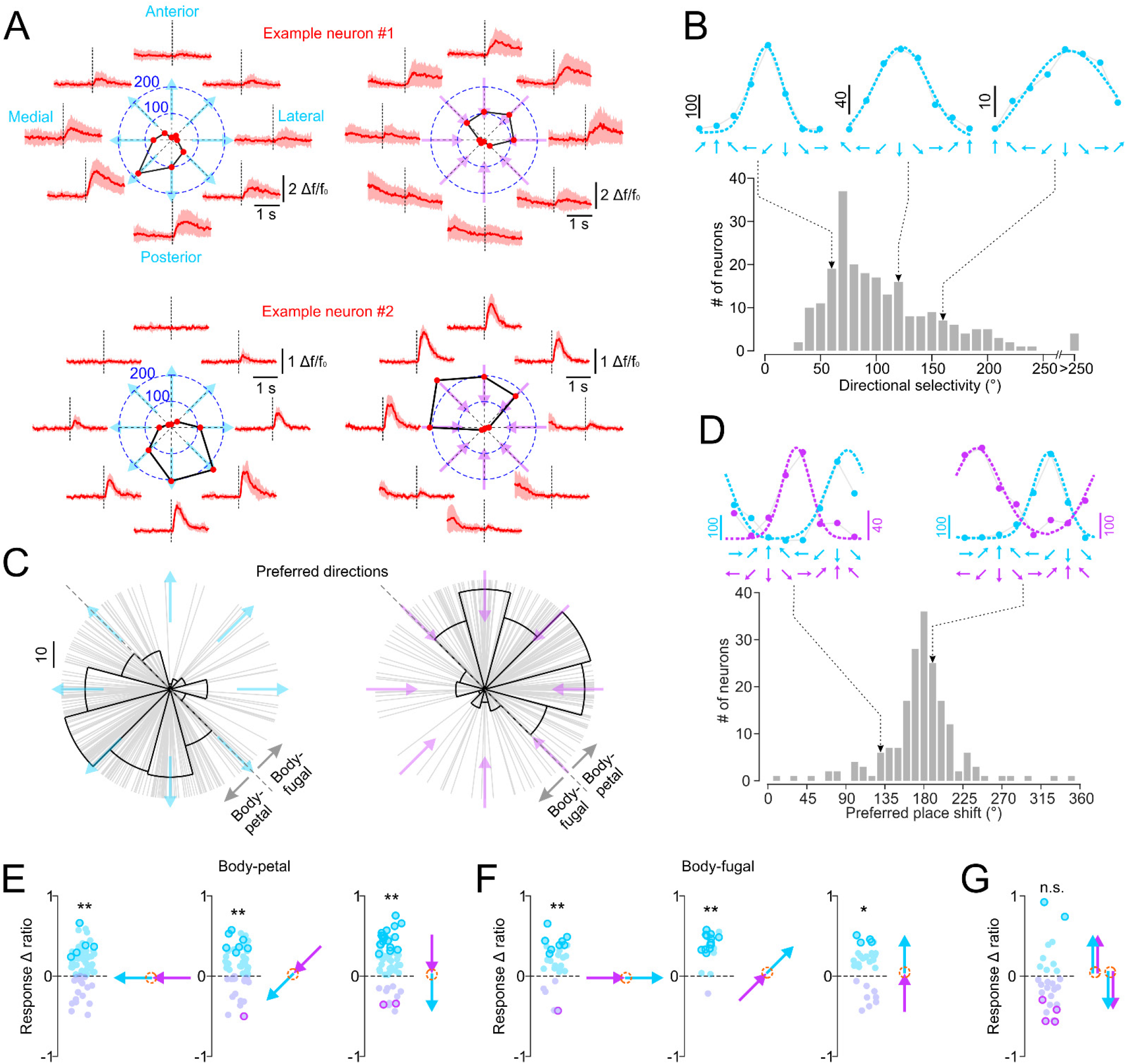
Selective tuning of fS1 neurons to the direction of passive forelimb movement revel a peripersonal representation of proprioception. **A**: Two example neurons with different preferred directions. Red traces: mean (± s.d.) responses to eight different directions of home-to-target movements (cyan arrows) and target-to-home movements (magenta arrows) with the same amplitude (7 mm) and velocity (2 cm/s). Polar plots: peak activity (deconvolved spike rate) as function of movement direction (red circles and numerical values refer to peak spike rate). Dotted lines: movement onset **B**: Top, Gaussian fits (dotted lines) to directionally tuned peak responses (deconvolved spike rate) of three example neurons. Bottom, distribution of directional selectivity (width of the Gaussian fits, measured in azimuth angles of the earth horizontal plane) for home-to-target movements (N=226 neurons, 18 mice). **C**: Distribution of preferred directions for home-to-target movements (cyan arrows, N=226 neurons) and target-to-home movements (magenta arrows, N=197 neurons) indicates a preferred representation of body-petal vs. body-fugal movements. **D:** Top, Gaussian fits to the directionally tuned responses (deconvolved spike rate) of two example neurons for home-to-target (cyan) and target-to-home (magenta) movements. Bottom, distribution of azimuth angle shifts (in the earth horizontal plane) in preferred spatial location between the two movement types (N=187 neurons). **E:** Response Δ ratios (see Methods for details) comparing peak neuronal activity for home-to-target vs. target-to-home movements with matched body-petal directions (orange circle: home position). Negative (magenta symbols) and positive (cyan symbols) values denote neurons with higher and lower activity for home-to-target movements, respectively. Bold symbols denote values significantly different from zero (p<0.01, two-sided t-test). **: p<0.01, *: p<0.05 (two-sided t-test) for the population mean. **F:** Same data and statistics as in E for matched body-fugal movements. **G:** Same data as in E comparing anterior and posterior movements with matched directions and start/end positions. n.s.: p=0.38.

The non-uniform distribution of preferred directions is consistent with body-fugal vs. body-petal coding of proprioception. The overrepresentation of body-petal directions might also suggest a preference for movements that bring the limb inside as opposed to outside the peripersonal space. A peripersonal representation implies that movements with matched directions should activate neurons differently if they start/end at different locations. To test this, we compared neuronal activations by home-to-target and target-to-home movements when their directions were matched (Fig. 6E,F). We observed that neuronal responses were significantly higher for body-petal movements that brought the limb closer to the body (Fig. 6E) and higher for body-fugal movements that brought the limb further away from the body (Fig. 6F). The higher activity cannot be explained by increased joint angle rotations, because home-to-target movements rotated the joints through the same absolute angles as target-to-home movements in opposite directions. For a subset of neurons, we systematically changed the starting home position so that we could compare home-to-target and target-to-home movements that are matched in both direction and start/end location. The difference was not significant in that case (Fig. 6G), suggesting a peripersonal representation of forelimb proprioception in fS1.

It follows that a muscle, joint or tendon input in primary afferents^38^ is elaborated along the ascending pathway into a perceptually relevant code in the cortex. In a new set of experiments, we tested whether this transformation results in categorically changing neural response sensitivity to movement kinematics. We measured how fS1 neurons are modulated by the amplitude and velocity of the passive movement stimuli in their preferred direction. Whereas primary and secondary muscle spindle afferents are linearly tuned to both the size and rate of change of muscle length^37^, we observed that the phasic responses of fS1 neurons are on average sensitive to amplitude (Fig. 7A,B) but that only a minority is significantly modulated by movement velocity (Fig. 7C,D). Movement size seems to be more relevant than its velocity for the encoding of peripersonal space and might thus explain this categorical difference between peripheral and cortical selectivity to proprioceptive stimuli.

**Figure 7.**
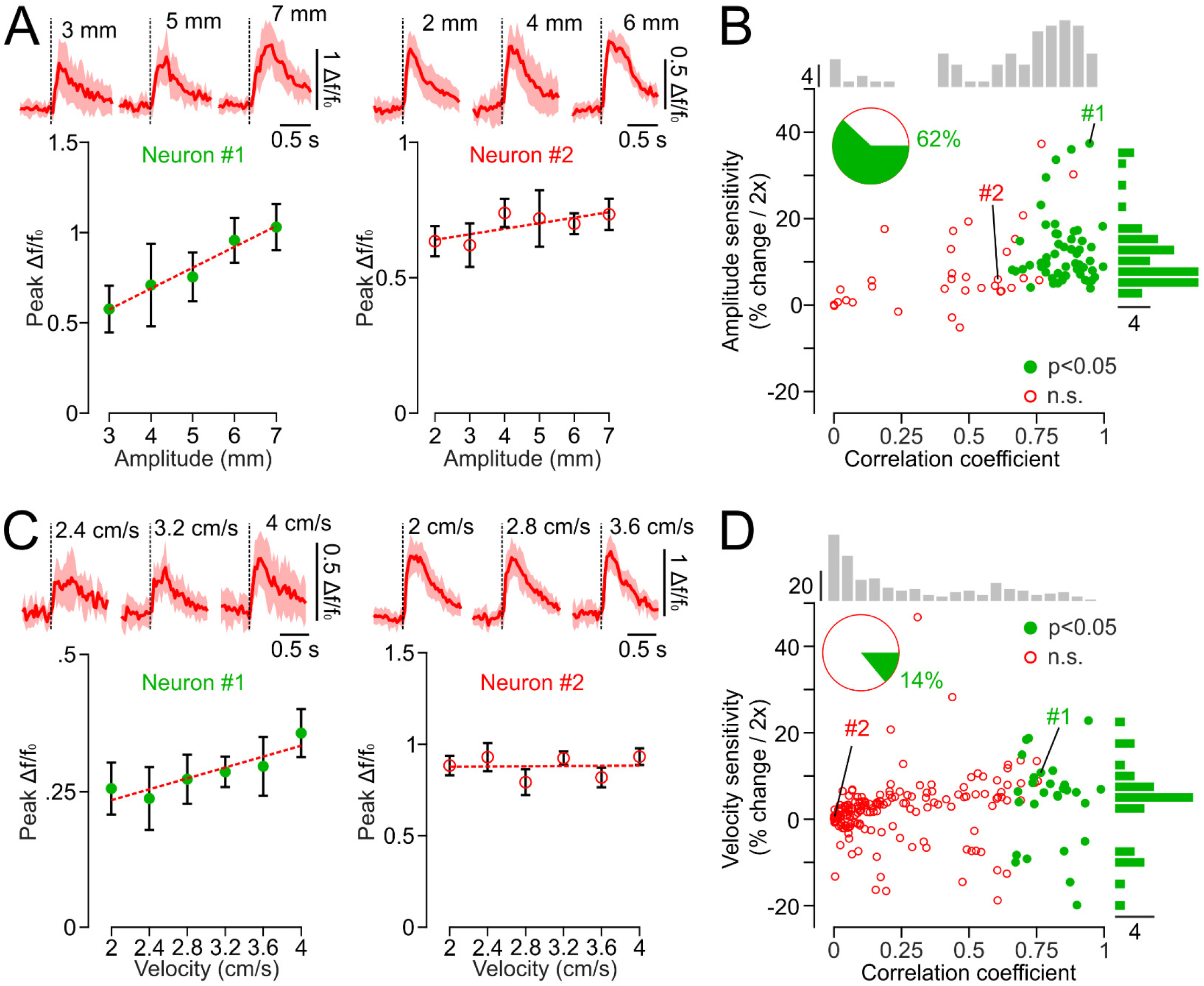
Amplitude and velocity tuning in fS1. **A:** Mean (± s.d.) peak responses of two example neurons tuned by (left) and insensitive (right) to displacement amplitude (dotted lines are linear regression fits). Top, *Δf/f_0_* mean (± s.d.) traces of the same neurons for three different amplitudes. **B:** Amplitude sensitivity (% change in peak response for a doubling of movement amplitude) as a function of correlation coefficient (peak response vs. movement amplitude) of neurons tested with varying amplitudes. Top histogram: distribution of correlation coefficients across all neurons. Right histogram: distribution of amplitude sensitivity for neurons with significant correlation with movement amplitude (N=86 neurons, 6 mice). The two example neurons in A are depicted with #1 and #2. **C, D:** Data analogous to that in A, B for neurons tested with varying velocities (N=197 neurons, 8 mice).

## Discussion

### The mouse proprioceptive cortex

Both proprioceptive (this study) and tactile^25,44^ neurons co-exist in mouse fS1 with seemingly little functional overlap (Fig. 5E). Optogenetic silencing of fS1 impairs the perception of proprioceptive stimuli (Fig. 3) and perceptual discrimination of vibrotactile frequencies also depends on fS1 since it follows the same computation rule as fS1 neuronal activity^25^. These findings indicate that, unlike primate areas 3a and 3b, proprioceptive and cutaneous somatosensation do not seem to be segregated in the mouse brain in terms of cortical territory. In that respect, fS1 is more similar to Brodmann’s area 2 that receives a combination of proprioceptive and tactile inputs^2^ and whose inactivation also evokes a proprioceptive deficit^45^.

Our results also show that the mouse proprioceptive cortex extends beyond fS1. It comprises the caudal forelimb motor cortex CFA and possibly a transitional zone TZ^33,35^ between the latter and fS1 (Fig. 3F). Indeed, responses to joint movements were qualitatively described in the rat fS1, TZ and CFA analogous areas^33^. This study reported that responses were most consistently found in TZ, which was also the only area with neurons responding to joint manipulation under anaesthesia. We found that the decrease in correct answers was the strongest and most consistently observed when the medial end of fS1 was silenced (Fig. 5E). Might this correspond to TZ and indicate the existence of a mouse homologue of the primate area 3a? The cytoarchitecture of 3a is markedly different from 3b in that it has an attenuated granular layer 4 (L4) and a thick layer 5 (L5). In the mouse cortex, it has recently been documented that the cytoarchitectural transition between fS1 and CFA is gradual^46^; from a cell sparse L5 in fS1 to a denser L5 in CFA and a progressive narrowing of L4 from fS1 to CFA, which is not completely agranular as classically described. Strikingly, a 3D analytical reconstruction revealed individual variations in the architectonic fS1-CFA boundary, which is evocative of the individual variability between animals in the location of area 3a with respect to the central sulcus^23^. We likewise found that the cortical locus of peak proprioceptive, but not tactile, activation was highly variable across mice (Fig. 2D). This variability might have occluded a clearer identification of a 3a homologue with our imaging and optogenetic silencing experiments.

### Proprioceptive representation of limb movement in mouse somatosensory cortex

A comparison between directional tuning curves for home-to-target and target-to-home movements (Fig. 6) revealed that most imaged neurons encode the direction of movement (a vector code) rather than hand location or limb posture (a position code). The non-uniform distribution of preferred directions showed a striking preference for movements that brought the limb closer to the mouse. We therefore propose that the cortical code specifies whether the limb’s endpoint is being displaced away from or towards a behaviorally relevant target (e.g. the body) rather than, for example, whether the elbow joint is being flexed or extended. To paraphrase Sherrington^47^, the body is therefore itself acting as a stimulus to its own limb movements.

Behavioral measurements suggest that this code guides decisions in a perceptual discrimination task (Fig. 3H, 4).

Non-uniform representations of preferred directions of limb movements have also been reported in the primate S1^6,48^, cerebellum^49^, cuneate nucleus^50^ and the cat dorsal spinocerebellar tract neurons^51^. The ansiotropy in the distribution was mostly bimodal along the flexion/extension axis of the limb (anteroposterior direction), suggesting an encoding of spatial information in a limb-based coordinate system^52^. In the mouse fS1, we however found a single mode along the body-fugal/petal axis (neurons strongly favor limb flexion and adduction over extension and abduction). Our findings thus support the idea that the proprioceptive cortex interfaces the limb with the body’s peripersonal space^53^, as opposed to being a transformation of a feedforward sensory map emerging from afferent innervation^54^. This idea is consistent with the observation that the topographic organization of area 3a in primates, and its homologue in other taxa, does not reflect innervation density but emerges and can reorganize as a result of the actual use of the limb in species-specific behaviors ^23^.

How is then this proprioceptive spatial direction signal generated in fS1? Afference from peripheral proprioceptors (e.g. muscle spindles) must be transformed along the ascending pathway to yield cortical responses that have “lost” their muscle or joint specificity. Accordingly, muscle length inputs to a cortical neuron could be continuously tuned by activity representing spatial information in somatosensory cortex^43,55^. Spatial activity could be acquired in fS1 based on direct connectivity from limbic structures^56,57^ or cortical areas interconnected with the hippocampal-entorhinal formation^58–60^. Encoding of space could in fact be a common feature of sensory cortical circuits^61,62^. The observed activity might actually not represent spatial information per se, but instead be a consequence of body simulations proposed to be the key functionality of the somatosensory cortex^54^. The proprioceptive code would thereby specify the movement of a particular body part with respect to another, such as the limb endpoint with respect to the trunk. It has been suggested that proprioceptive coding in relation to the body or its peripersonal space might be computed by the reciprocally connected circuit between somatosensory cortical regions and the pulvinar^63,64^, or its homologue in the mouse thalamus^65^.

Alternatively, this “high level” code could be inherited, at least in part, from second order dorsal column neurons that represent limb proprioception more in terms of global parameters than joint angles or muscle lengths^52,66,67^. A simulation study suggests that such signals can theoretically also arise from randomly weighted muscle spindle inputs^68^. Because second order neurons receive direct excitatory and inhibitory inputs from corticofugal axons^21,69^ it remains unclear whether these signals are first computed in fS1, at early stages of the pathway, reflect musculoskeletal geometry or a combination thereof. It is generally difficult to discern how much peripheral inputs actually need to be transformed to explain a neural proprioceptive code. Spindle afferents from passive muscles signal more than just information related to stretch^70^. Many muscles are biarticular (span two joints) and biomechanical constraints between different limb segments and joints can result in signals related to global limb representations that are not necessarily indicative of central neural processing^52^. Future experiments using multi-site imaging should more directly compare signals at all levels of the ascending pathway to identify key computational transformations of the proprioceptive code.

### Implication for neuroprosthetics

To fully replace a paralyzed or lost limb, a neural prosthesis must be bidirectional: as it decodes motor signals, it must simultaneously deliver sensory signals to mimic proprioceptive feedback. One strategy is to stimulate the somatosensory cortex to provide a proprioceptive-like sensation of the prosthetic movement^71,72^. A crucial question is what kinematic features of the movement should the stimulation paradigm be based on? Our findings imply that stimulation patterns should be correlated with a movement direction vector of the hand (or its spatial trajectory) rather than with a combination of joint angles.

In agreement with our results, discharge rates of neurons in primate S1 to static arm postures show less variability when plotted against parameters describing spatial hand location than orientation angles in joint space^48^. Psychophysical data in humans show that for passive arm displacements, the perception of arm endpoint^73^ and the orientation of the limb relative to gravity^74^ is more precise than the perception of joint angles. Similarly, illusory movements evoked by stimulation of afferents from groups of muscles are not perceived in terms of neuroprosthetic control.

We argue that a stimulation paradigm based on the neural representation of passive proprioception is better suited for neuroprosthetic movement restoration. If the aim is to evoke a proprioceptive-like percept, then sensory ex-afference is perceptually more salient than sensory re-afference resulting from active movements^8^. Muscles in paralyzed or non-existent limbs do not contract and decoded motor activity bypasses a large part of the descending motor circuitry. Therefore, engaging the perceptual instead of the motor proprioceptive pathway^3^ seems to be more relevant for muscle length or joint angle changes but in terms of the displacement of the limb’s endpoint along a given spatial trajectory^8,75^. On the contrary, proprioceptive responses of neurons in primate area 2 during reaching movements are better explained by a model based on muscle lengths^76^ or whole-arm kinematics^3^ than by a hand-only model. Joint angles were also more precisely estimated in active versus passive movements^73^. It thus seems that passive proprioceptive afference is preferentially encoded in the cortex and perceived in terms of spatial endpoint kinematics, but that during active behavior the contribution from motor commands (i.e. efference copies)^5^ and the influence of the fusimotor drive^9,10^ increases the complexity of the cortical code.

## Methods

### Mice

For Ca^2+^ imaging of cortical neurons we used 16 male C57BL/6 mice (Charles River Laboratory) and 3 male Thy1-GCaMP6f-GP5.17 mice (Jackson laboratory; stock no. 025393). For optogenetic silencing experiments we used 1 male and 3 female VGAT-ChR2-eYFP mice (Jackson laboratory; stock no. 014548). For wide-field Ca^2+^ imaging we used 4 female and 2 male double transgenic mice obtained by crossing Rasgrf2-dCre (Jackson laboratory; stock no. 022864) with Ai148 mice (Jackson laboratory; stock no. 030328), a TIGRE2.0 Cre-dependent GCaMP6f reporter line. For anatomical tracing experiments we used 3 double transgenic mice generated by crossing PV-Cre (Jackson laboratory; stock no. 017320) with Ai32 mice (Jackson laboratory; stock no. 024109), a Cre-dependent ChR2/EYFP reporter line, to trace proprioceptive afferents and 3 C57BL/6 mice (Charles River Laboratory) to trace cuneo-thalamic projections. All were 6 to 12 weeks old at the start of experiments. Mice were housed in an animal facility in groups of maximum five per cage, maintained on a 12h/12h light/dark cycle and placed on a water restriction regime of 1 ml/day during experiments. All procedures were approved by and complied with the guidelines of the Fribourg Cantonal Commission for Animal Experimentation.

### Muscle and thalamic virus injections

Virus injections were made in adult (6 to 8 week old) mice anesthetized with 2% isoflurane and immobilized on a motorized stereotaxic frame. For muscle injections, the skin of the right forelimb of the mouse was opened and the triceps and biceps were exposed. AAV9/2-CAG-dlox-tdTomato (Zurich Viral Vector Facility, v167-9, stock titer 5.5×10^12^ vg/ml) was injected in the muscles (total volume of 1.5 μL) with a glass micropipette. The skin was sutured and the brain perfused 4 weeks later. For thalamic injections, the skull was exposed and AAV-retro/2-CAG-EGFP (Zurich Viral Vector Facility, v24-retro, stock titer 5×10^12^ vg/ml) was targeted to VPL (−1.9 mm posterior, −2.0 mm lateral, −3.7 mm deep; 60 nL volume) and AAV-retro/2-shortCAG-tdTomato (Zurich Viral Vector Facility, v131-retro, stock titer 7.1×10^12^ vg/ml) was targeted to PO (−1.9 mm posterior; - 1.2 mm lateral; −3.3 mm deep; 100 nL volume) through a small craniotomy with glass micropipettes (tip diameter 30 to 40 μm). The skin was sutured and the brain perfused 2 weeks later.

### Immunohistochemistry

Mice were transcardially perfused with 4% ice cold paraformaldehyde (PFA, in 0.1 M sodium phosphate buffer, pH 7.4). Cervical DRGs and brain were immediately dissected and post-fixed for 2.5 h with 4% PFA on ice. Post-fixed tissue was washed (3 x 10 min) with 0.1 M sodium phosphate buffer (pH 7.4) and then incubated in 30% sucrose (in phosphate buffered saline, PBS) overnight at 4°C for cryoprotection. Cryoprotected tissue was cut at 16 μm or 40 μm (DRGs or brain, respectively) on a cryostat (HM525 NX, Thermo Scientific), mounted on Superfrost Plus glass slides and then incubated with the respective combinations of primary antibodies (Rabbit anti-GFP, 1:1000, A-6455, Thermo Fisher; Goat anti-tdTomato, 1:1000, AB8181-200, SICGEN) in 1% donkey serum in PBS over-night at 4°C. After washes in PBS (3 x 10 min), sections were incubated with the respective secondary antibodies (Alexa Fluor 488 Donkey anti-Rabbit, 1:500, AB_2313584, Jackson Immuno Research; Cy3 Donkey anti-Goat, 1:500, AB_2307351, Jackon Immuno Research) for 2 h at room temperature and rinsed in PBS (3 x 10 min), before mounting with coverslips and fluorescent Dako Mounting Medium (Agilent Technologies).

### Surgical procedures for two-photon imaging experiments

Surgeries were performed under isoflurane anesthesia (1.5 to 2% in 1.5 L/min O_2_). We administered additional analgesic (0.1 mg/kg buprenorphine intramuscular (i.m.)), local anaesthetic (75 μl 1% lidocaine subcutaneous (s.c.) under the scalp) and anti-inflammatory drugs (2.5 mg/kg dexamethasone i.m. and 5 mg/kg carprofen s.c.). Mice were fixed in a stereotaxic frame and rested on a heating pad (37°C). An incision was made over the midline between the ears and eyes to expose the scalp. To allow for head fixation during experiments, a titanium head frame was fixed on the skull with a cyanoacrylate adhesive (ergo 5011, IBZ industrie) and clear dental acrylic (Paladur, Kulzer GmbH). We made a craniotomy centered over the left forelimb somatosensory cortex (fS1) and performed five viral injections at stereotaxic coordinates −2.25 mm lateral and from −0.25 mm to 0.75 mm in 0.25 mm steps anterior to bregma (based on localization of fS1 in mice with intrinsic signal imaging ^25^) using pulled and beveled (≈25 μm tip diameter) glass pipettes (Wiretroll II, Drummond Scientific). We injected AAV9/2-hSyn1-jGCaMP7f (Zurich Viral Vector Facility, v292-9, stock titer 4.4×10^12^ vg/ml) in 12 C57BL/6 mice and AAV9/2-hSyn1-jGCaMP8m (Zurich Viral Vector Facility, v623-9, stock titer 6.4×10^12^ vg/ml) in 4 C57BL/6 mice (1:10 dilution with 0.2% FastGreen in sterile saline) at a depth of 350 μm, 30 to 60 nl per site at a rate of 20 nl/min. After rinsing the cortical surface with dextamethasone (0.01 ml of a 4 mg/ml solution) we covered the craniotomy with a cranial window. The window consisted of two hand-cut glass coverslips (150 μm) glued together with optical adhesive (NOA 61, Norland). The lower one, matching the shape of the craniotomy, was placed on the cortical surface and the top one, cut to 1 mm larger than the craniotomy, was fixed to the skull with cyanoacrylate glue and dental acrylic. Experiments typically began 14 days after surgery. The same surgery but without viral injections was performed in the Thy1-GCaMP6f-GP5.17 mice.

### Surgical procedures for cortical silencing experiments

Under the same anesthesia protocol, VGAT-ChR2-eYFP mice were implanted with a titanium head frame as above. We made a transparent skull preparation for transcranial optical access^31,77,78^. All periosteum was removed from the skull surface and the area thoroughly cleaned. The skull surface was homogenously covered with a thin layer of transparent dental acrylic (Paladur, Kulzer GmbH). After curing, a drop of cyanoacrylate adhesive (ergo 5011, IBZ industrie) was spread on the coated surface and made the skull transparent.

### Surgical procedures for wide-field imaging experiments

The same transparent skull surgical procedure was performed as above. After recovery, to induce recombinase activity of dCre we injected Rasgrf2-dCre;Ai148 mice intraperitoneally with trimethoprim (Sigma-Aldrich T7883) at 0.25 mg/g body weight per day for 3 days. Trimethoprim was dissolved in DMSO (Sigma-Aldrich 34869) at 100 mg/ml. The stock solution was diluted with 0.9% NaCl immediately prior to injection.

### Robotic manipulandum

We custom-built a robotic manipulandum based on the planar 2 DOF pantograph design^79–81^. The robot consists of four CNC machined aluminum arm linkages connected to each other at three joints (Supplementary Fig. 1A) using miniature ball bearings (Reely MR 52 ZZ, 2 mm Ø). A handle (steel rod, 2 mm Ø, with the tip rounded for comfortable grasping) is mounted at the endpoint joint. The mechanism is mounted on and actuated by two DC motors (DCX22L EB SL 9V, Maxon Motors) with integrated optical rotary encoders (ENX 16 RIO, 32768 counts/turn, Maxon Motors). A 1:16 reduction gear (GPX22 A, Maxon Motors) is mounted on each unit to maximize position stability during actuation (i.e. uniformly counteract the impedance of the mouse limb, Supplementary Fig. 1C,D) and increase angular positioning resolution. The motors are operated in position mode with the EPOS2 24/5 positioning PID controllers (Maxon, 1 kHz sample rate) and interfaced via USB with Matlab using EPOS2 libraries. Transformations between angular coordinates of the motors and planar Cartesian coordinates of the manipulandum’s endpoint are coded in Matlab by computing the forward and inverse kinematics of the linkage structure^79^. The angular position of each motor was read via USB and used to compute and monitor online the instantaneous position of the manipulandum at a rate of 100 Hz. In parallel, we recorded the position at a 1 kHz sampling rate with a custom-built circuit. The quadrature signals from the optical encoders were decoded using the hardware quad decoders of Arduino DUE and the 16-bit digital signals at its output transformed to analog signals (AD669ANZ, Analog devices). The analog signals were sampled at 1 kHz (NI PCIe-6321, National Instruments) and logged to disk.

### Behavioral procedures

All behavior was controlled and measured with real-time protocols using the Bpod State Machine r1 system (Sanworks) interfaced with Matlab. We created a Matlab object as a plugin to the Bpod code in order to control the robotic manipulandum from within a Bpod protocol.

#### Passive forelimb movement task

Mice sat head fixed inside a tube and trained to hold the robotic manipulandum handle with their right paw. The home position (i.e. the forelimb endpoint) was located approximately 17 mm below, 2.5 mm lateral and 10 mm posterior to the mouse snout. Contact with the handle was detected with a capacitive sensor (MPR121, Adafruit interfaced with an Arduino Nano Every board). Each trial began with a pre-stimulus baseline requiring 2 s of continuous holding. A release resulted in resetting the 2 s wait period. The manipulandum was then displaced radially from its home position to a target position in one of 8 co-planar cardinal directions with a trapezoidal velocity profile (3 cm/s). The movement amplitude was between 5 and 8 mm. After a random 1 to 2 s holding period at target position, the manipulandum returned to its home position followed by a second 1 to 2 s random holding period (Supplementary Movie 1). Releasing the handle at any time during the trial resulted in a punishment (air puff to the face) and an aborted trial. Continuous holding resulted in a correct trial and a water droplet reward (Fig. 2B). In the amplitude and velocity experiments (Fig. 7), between 1 and 4 directions were tested simultaneously. At most seven and at least five different amplitudes between 2 mm and 8 mm (at a fixed velocity) and six different velocities between 2 and 4 cm/s (at a fixed amplitude) were tested.

#### Perceptual discrimination task

Mice had to perceptually discriminate between two directions of passive forelimb movement with a directional lick toward one of two reward spouts (Supplementary Movie 2). Each trial started with a 2 s pre-stimulus period (as above) requiring continuous holding and no licking of the reward spouts. During the subsequent stimulus period, the manipulandum passively displaced the right mouse forelimb either laterally (i.e. abduction) or medially (i.e. adduction), stayed at the target position for 400 ms and returned home. An auditory mask (white noise sampled at 50 kHz to cover the hearing range of the mouse) was played on a loudspeaker during the stimulus to mask the sound of the motors. We tested displacement sizes between 1 and 4 mm at a 2 cm/s velocity. The optogenetic silencing results are based on either 4 or 3 mm displacements (Fig. 3). The stimulus was followed by a delay period, an auditory go cue and an answer period (Fig. 3B). During the answer period (limited to 2 s), if the mouse licked the correct water spout (right for lateral and left for medial for 1 mouse and the reversed contingency for 3 other mice) he received a water droplet at that spout or an air puff for the incorrect licking direction. For analysis, we standardized the stimulus/answer rule to the normal contingency. Releasing the handle during stimulation, licking during the stimulus or delay periods and not answering resulted in an aborted trial and a 4 s timeout. Holding and licking was detected with a capacitive sensor (MPR121, Adafruit). To minimize a directional licking bias, the probability of a medial trial *(P_med_)* was determined in real-time as a function of the measured bias during the last 10 non-aborted trials. The bias value was calculated as the difference in the fraction of correct responses between medial and lateral trials. *P_med_* was calculated at the start of each trial according to the double sigmoidal function:

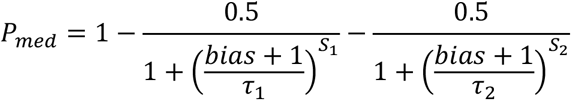

where the inflection slopes *S_1_* and *S_2_* at the chosen inflection points τ_1_=-0.5 and τ_2_=0.5 were set to 30 and 12 respectively.

In the first 5 to 7 days of training, the reward was automatically delivered at the correct spout during the go-cue while maintaining all trial abort rules. This allowed mice to first learn the abort rules as well as the stimulus/response association. In expert mice (>75% correct performance), probe stimuli (posterior and lateral movements, Fig. 3G,H) were tested in 10 sessions/mouse and occurred randomly on 15% of trials. They were matched in amplitude and velocity to the trained stimuli (medial and lateral movements).

#### Tactile stimulation

Tactile stimuli (Fig. 5E,F) were automatized indentation of the paw’s glabrous skin using a custom-built device. A nylon bristle (0.35 mm diameter) was mounted on a push-pull solenoid (Adafruit 412) actuated by relaying a 12 V signal from a high current source (custom circuit) with a TTL pulse from Bpod. Foam material was added to the solenoid base to limit its full travel and thereby mask sound. The displaced bristle traveled through the center of a paw holder (3D printed) and evoked a ≈1 mm skin indentation lasting 500 ms. Successive stimuli were separated by at least 3 s, occurred randomly and only when the mouse had its paw placed on the holder.

#### Vibrotactile stimulation

Vibrotactile stimuli (Fig. 2C) were 100 Hz vibrations lasting 0.5 s transmitted through a handle (steel rod, 2 mm Ø) to the mouse paw. The handle was mounted on a galvanometric actuator (PT-A40, Phenix Technology) controlled by analog signals from the Bpod analog output module (Sanworks). The trial structure and all other experimental details were as for the passive forelimb movement task.

#### Nerve block

Mice were briefly anaesthetized with isoflurane (3%). Neural transmission from the forepaw was blocked with a single 10 μL injection of lidocaine (1%) in the palm (s.c.). Mice were subsequently head-fixed under the two-photon microscope and allowed to recover from anaesthesia for ≈10 min before starting the experiment. The imaged responses were compared to those of the same neurons obtained pre-injection (Fig. 5F).

### Two-photon microscopy

Ca^2+^ imaging in the mouse cortex was performed with a custom built two-photon microscope based on an open source design (MIMMS 2.0, janelia.org/open-science) and controlled with Scanimage 5.7_R1 software (Vidrio Technologies) and National Instrument electronics. The Ti:Sapphire excitation laser (Tiberius, Thorlabs) was tuned to 930 nm and focused with a 16x 0.8 NA objective (Nikon) below the cortical surface. The laser power (typically 25 mW measured at the objective) was modulated with pockels cell (350-80-LA-02, Conoptics) and calibrated with a Ge photodetector (DET50B2, Thorlabs). A 550 μm by 550 μm area of cortex was scanned at ≈30 frames/s using a resonant-galvo scanning system (CRS8K/6215H, CRS/671-HP Cambridge Technologies). Emitted fluorescence was detected with GaAsP photomultiplier tubes (PMT2101, Thorlabs) and the acquired 512 x 512 pixel images written in 16 bit format to disk. Behavioral event (trial start and stimulus onset) TTL pulses issued by the Bpod State Machine were received as auxiliary inputs to the Scanimage electronics and their timestamps saved in the headers of the acquired images. The timestamps were used to temporally align neuronal data to behavioral events.

### Optogenetic silencing

Cortical silencing was achieved by optogenetic activation of GABAergic cortical neurons (i.e. indirect inhibition of excitatory neurons) in VGAT-ChR2-eYFP mice through the clear skull preparation using a 473 nm laser (Obis LX FP 473, Coherent) operated in analog control mode. This silencing method is shown to be more effective than direct inactivation of excitatory cells using inhibitory opsins^82^. The optical fiber from the laser was inserted into an aspheric collimator (CFC11A-A, Thorlabs) and the resulting free space beam aimed at cortical coordinates with a pair of galvanometric scanning mirrors (PT-A40, Phenix Technology). The laser beam was focused on the cortical surface with an achromatic doublet lens (AC254-100-A, Thorlabs) and gated with a shutter (SHB05T, Thorlabs). The laser power and position of scanning mirrors were controlled by analog signals from the Bpod analog output module (Sanworks). The laser power modulation signal was a 40 Hz sinusoid of duration equal to the stimulation period. The last 100 ms of the signal were ramped down linearly. The mean power of the stimulation signal used in our experiments was 1.5 mW (measured at the cortical surface). Scanner controller voltages corresponding to the coordinates of the silenced regions (Fig. 3D,F) relative to bregma (fS1: −2.25 mm lateral, 0.25 mm anterior; hS1: −1.75 mm, −0.75 mm; ALM: −1.5 mm, 2.5 mm; ipsi-fS1: 2.25, mm, 0.25 mm; wS1: −3.5 mm, −1.5 mm; CFA: −1.4 mm, 250 mm) were calculated using a calibration head frame and the reference (0,0) coordinate was aligned to bregma at the start of each session. The coordinates for each mouse were registered to the Allen Mouse Brain Atlas (Supplementary Fig. 2) using the same procedure as for wide-field data analysis (see below). The line stimuli (Fig. 3F) were 1 mm long, centered on 0, −0.5, −1, −1.5, −2, −2.25, −2.5, −3 and −3.5 mm lateral to bregma, and produced by a 40 Hz triangular wave oscillation of the scanning mirror. Three targets were inactivated per session on 50% of the trials. On the remaining 50%, the laser was aimed at a control site outside of the cortical surface (posterior end of the head frame). The inactivation trials were therefore not visually cued as the blue light stimulus was present on every trial. 5 inactivation sessions were performed per mouse and target.

### Wide-field imaging

Wide-field Ca^2+^ imaging of the mouse cortex (Fig. 2) was performed with a custom built fluorescence macroscope^83^. Top and bottom objectives (50 mm f/1.2 Nikon camera lenses; bottom lens inverted) were mounted on a 60 mm fluorescence filter cube (DFM2/M, Thorlabs) housing a dichroic mirror (495 nm beamsplitter, T 495 LPXR, AHF). Blue 470 nm and violet 405 nm LED light (M470L5 and M405L4, Thorlabs) was collimated, diffused (ACL2520U-DG6-A, Thorlabs), passed through excitation filters (MF469-35, Thorlabs; #65-133 Edmund Optics), combined with a beamsplitter (435 nm dichroic filter, #87-063) mounted inside a 30 mm filter cube (DFM1/M, Thorlabs) and coupled into the 60 mm cube with a third lens (50 mm f/1.2, Nikon). Emitted green light passed through both objectives and an emission filter (525/45 nm, Edmund Optics) allowing images to be acquired with a sCMOS camera (ORCA Flash 4.0 LT+, Hamamatsu) after focusing ~100 μm below blood vessels. The total power of excitation light on the surface of the brain was measured to be below 5 mW. Images were acquired at 40 fps and 512 by 512 pixel resolution with alternating 470 nm and 405 nm illumination controlled with a microcontroller (Arduino UNO).

### X-ray assisted 3D joint tracking

The corpse of an adult mouse (≈25 g) was head fixed and its right forelimb stuck to the manipulandum endpoint using the same apparatus configuration as in the experimental condition. The apparatus was placed in a C arm fluoroscope (Philips BV 25) and the forelimb displaced in succession to each point of the planar workspace (as defined in Supplementary Fig. 1B). The acquisition (Matrox Solios eCL-B frame grabber) of the detected x-ray images (Dexela 1207 flat panel ray detector) were triggered by TTL pulses (NI PCIe-6321, National Instruments) from the PC controlling the manipulandum’s position (see above). The acquisition was repeated at two orientations relative to the source/detector axis (side and front views in Fig. 3A). Independently, we acquired video images at the same manipulandum positions after surgically removing the skin and adipose tissue of the mouse’s forelimb by two cameras simultaneously (Basler dart USB 3.0, daA1280-54um, 1280 x 960 resolution with a 8 mm Evetar IR lens). The two cameras were pointed at the skinned forelimb from two different orientations in the horizontal plane and externally triggered by the same TTL pulses. A side-by-side comparison of the X-ray and video images allowed us to precisely hand score the locations of the endpoint and 4 joints (scapulothoracic, glenohumeral, elbow and wrist) on the limb musculature of each stereo image pair using a custom graphic user interface programmed in Matlab. Using a checkerboard pattern and the Matlab *Stereo Camera Calibrator App* we obtained the stereo calibration parameters of our stereo camera configuration. We subsequently calculated the world 3D positions of each joint (with the optical center of camera 1 as the origin) using the *triangulate* function in Matlab by passing the stereo camera coordinates and calibration parameters as inputs. To transform the 3D positions from camera into manipulandum coordinates (where the origin is the center of the left motor as shown in Supplementary Fig. 1A,B), we first fit a geometric transformation based on rotation, scaling and translation (without reflection and shearing) between the manipulandum positions (in manipulandum coordinates) and the tracked positions of the endpoint (in camera coordinates) using the *fitgeotrans* Matlab function. The fitted transformation was then used to transform the camera into manipulandum coordinates with the *transformPointsForward* function. Prior to the transformation, the values were converted from mm to cm, the Y and Z axes swapped and the Z axis inverted.

We calculated the humerus abduction/adduction angle (Fig. 4A) from the 3D joint coordinates (Supplementary Movie 3) for each position on the planar workspace. The calculation of the angle is graphically defined in Fig. 4A (above the color map): humerus abduction/adduction is the azimuth angle between the -Z axis vector and the vector of the humerus (link defined by points 2 and 3) projected on the XZ plane.

### Two-photon data processing

#### Motion correction

A custom MATLAB registration script was used to correct for vertical and horizontal image movements. Each acquired image was aligned to a baseline average image recorded at the start of each session. We computed the crosscorrelation between each image and the template by multiplying the two-dimensional discrete Fourier transform of one with the complex conjugate of the Fourier transform of the other and taking the inverse Fourier transform of the product. The X and Y location of the peak cross-correlation value gave the vertical and horizontal shift, respectively. 10% of each image was cropped at the boundaries before carrying out the computation.

#### Region of interest and Ca^2+^ activity generation

Using the session mean and variance images, soma centers of active neurons with clearly identifiable morphologies were manually initialized. Regions of interest of individual neurons and background were then identified as spatial footprints using the constrained nonnegative matrix factorization method^84^ from the CaImAn Matlab toolbox (github.com/flatironinstitute/CaImAn-MATLAB). The time-varying calcium activity of each neuron (i.e. its spatial footprint) and their time-varying baseline fluorescence was subsequently extracted from the acquired images and used to compute *Δf/f_0_* traces used for analysis.

##### Spike rate deconvolution

Spike rate was inferred from the *Δf/f_0_* traces using the OASIS deconvolution algorithm^85^ of the Suite 2P toolbox (github.com/cortex-lab/Suite2P) with a 0.8 s sensor timescale. Spike rate density was calculated by convolution of the inferred spikes with a Gaussian kernel (0.1 s width, 1 kHz sampling rate) and multiplying the result with the sampling rate.

### Wide-field data processing

All analysis was performed with custom written routines in Matlab (Mathworks). The frame timestamps of images acquired with blue and violet light excitation were interpolated to the same regular 20 Hz time points and stimulus aligned averages (>80 trials/session) were computed for each channel. The violet channel was regressed onto the blue channel for each pixel. The violet channel was then scaled with the obtained regression coefficients and subtracted from the blue channel to correct for hemodynamic signals and other non-calcium dependent artifacts. The corrected signal was *Δf/f_0_* normalized for each pixel by taking the mean over the 1 s preceding stimulus onset as baseline and temporally smoothed with a Savitzky-Golay filter (width = 450 ms, order = 2). Frames with peak activation were analyzed for each mouse and occurred on average 250 ms after stimulus onset. We registered each frame to the 2D top projection of the Allen Mouse Brain Atlas (mouse.brain-map.org) using the bregma, lambda, anterolateral tips of the left and right parietal bones and the median point between the two frontal poles as reference points^86^. The registration was performed for each mouse using the affine transformation *(fitgeotrans* and *imwarp* Matlab functions).

### Data analysis

#### Stimulus evoked responses

The stimulus evoked *Δf/f_0_* (or inferred spike rate) response was defined as the difference between the maximum post stimulus value (in the 0 to 1.5 s interval relative to onset) and the mean value pre-stimulus value (between 0 s and −0.75 s relative to onset). Significant responses were identified with a randomization test at significance level p<0.01. Specifically, the response calculation for each neuron was repeated 1999 times for randomly shifted stimulus onset times across the neuron’s activity trace of the session. The 1999 calculated chance measures were compared to the non-randomized response and the latter was deemed significant if it was more extreme than the upper 99^th^ percentile of the chance values distribution.

#### Directional selectivity

Directional selectivity was tested by fitting a Gaussian function 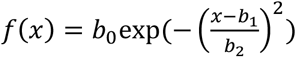 to the neuron’s stimulus-evoked responses (inferred spike rate) in the 8 tested directions (*fit* function in Matlab). The directional responses were first shifted circularly to center the data on the direction with the maximum response. A neuron was deemed to be directionally selective if the 95% confidence intervals of the *b_0_* and *b_2_* fitted parameters did not include zero. The neuron’s preferred direction and directional selectivity were defined by the *b_1_* and *b_2_* fitted parameters, respectively. The non-uniformity of preferred direction distribution was tested using the Rayleigh test (Circular Statistics Toolbox^87^, Matlab)

#### Comparison of responses for different starting positions

To test for a representation of peripersonal space we compared how neuronal responses differ between stimuli with matched direction vectors but different start/end positions (Fig. 6E-G). The response Δ ratio was computed for each neuron as:

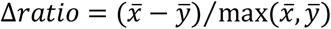

Where 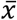 and *ȳ* are the average peak responses to home-to-target and target-to-home movements, respectively. For each comparison, this analysis included all neurons with significant responses (see *Stimulus evoked responses* paragraph above) for both movement types regardless of their preferred direction.

#### Behavioral data analysis

In the perceptual discrimination task, the Δ% correct (Fig. 3D,F) for each inactivation site was defined as the drop in % of correct responses compared to the control site. Data from 20 sessions (5 sessions/mouse) were pooled for each inactivation site.

#### Psychometric curve fitting

In the perceptual discrimination task, we analyzed the fraction of right lick answers as a function of directional displacement amplitude (Fig. 3C). The data was fit with a sigmoid function (i.e. a cumulative Gaussian, including the lapse rate and guess rate parameters) using the *psignfit* Matlab toolbox ^88^. Each mouse performed at least 36 and at most 90 trials per tested amplitude.

#### Statistics

Measurements for any one experiment were made from different animals/neurons and no neuron was measured repeatedly. The normality assumption was tested with the Kolmogorov-Smirnov test. Non-parametric tests were used when the normality assumption was not met.

## Data availability

The data generated and/or analysed during the current study are available from the corresponding authors upon reasonable request.

## Code availability

The Matlab code for generating, collecting and analyzing data in the current study is available from the corresponding authors upon reasonable request.

## Supplementary material

**Supplementary Figure 1.**
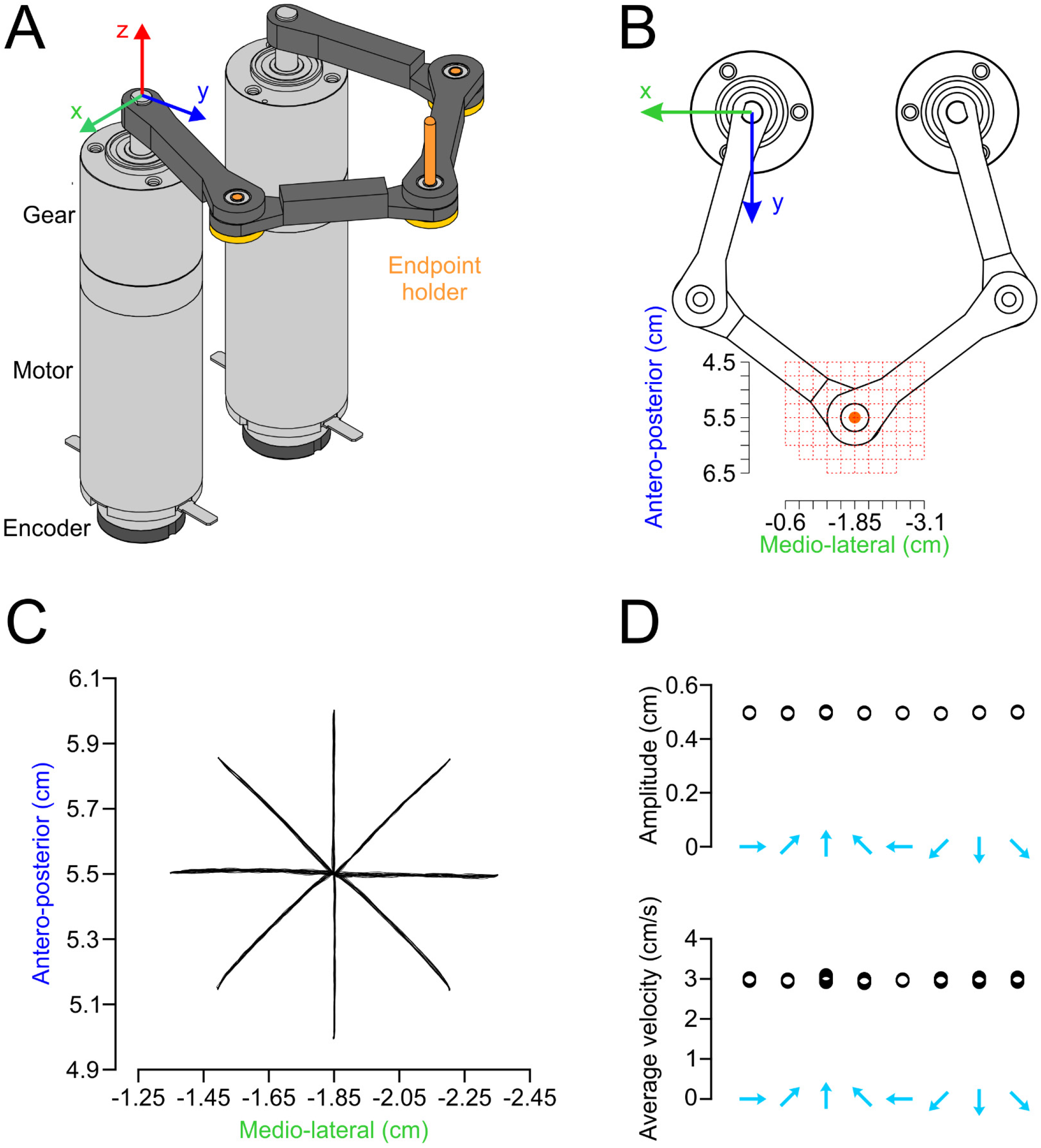
Movement kinematics of the robotic manipulandum. **A**: CAD model of the robotic manipulandum showing the origin of the Cartesian coordinates. **B**: Top view of the manipulandum showing the workspace of forelimb movements (red grid). **C**: Superimposed 2D trajectories of the manipulandum’s endpoint (raw unfiltered measurements with the optical encoders and sampled at 1 kHz, see Methods) for movements in the eight tested directions in an example session (17 to 23 trials per direction, 0.5 cm amplitude and 3 cm /s velocity). **D**: Measured 2D amplitudes and average velocities of the individual movements in A were highly consistent across the tested directions.

**Supplementary Figure 2.**
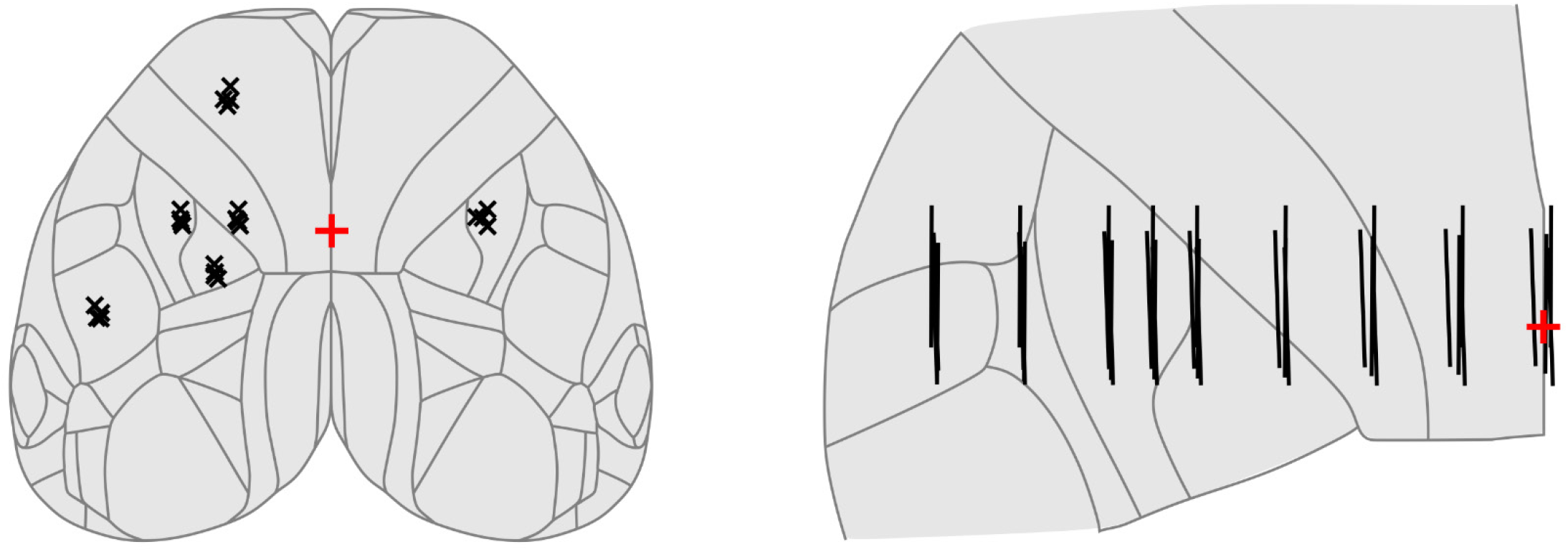
Optogenetic stimulation sites. Stimulation coordinates of the optogenetic stimuli (Fig. 3D, F) registered to the surface projection of the Allen Mouse Brain Atlas indicate consistent targeting of the same areas across the 4 mice.

**Supplementary Figure 3.**
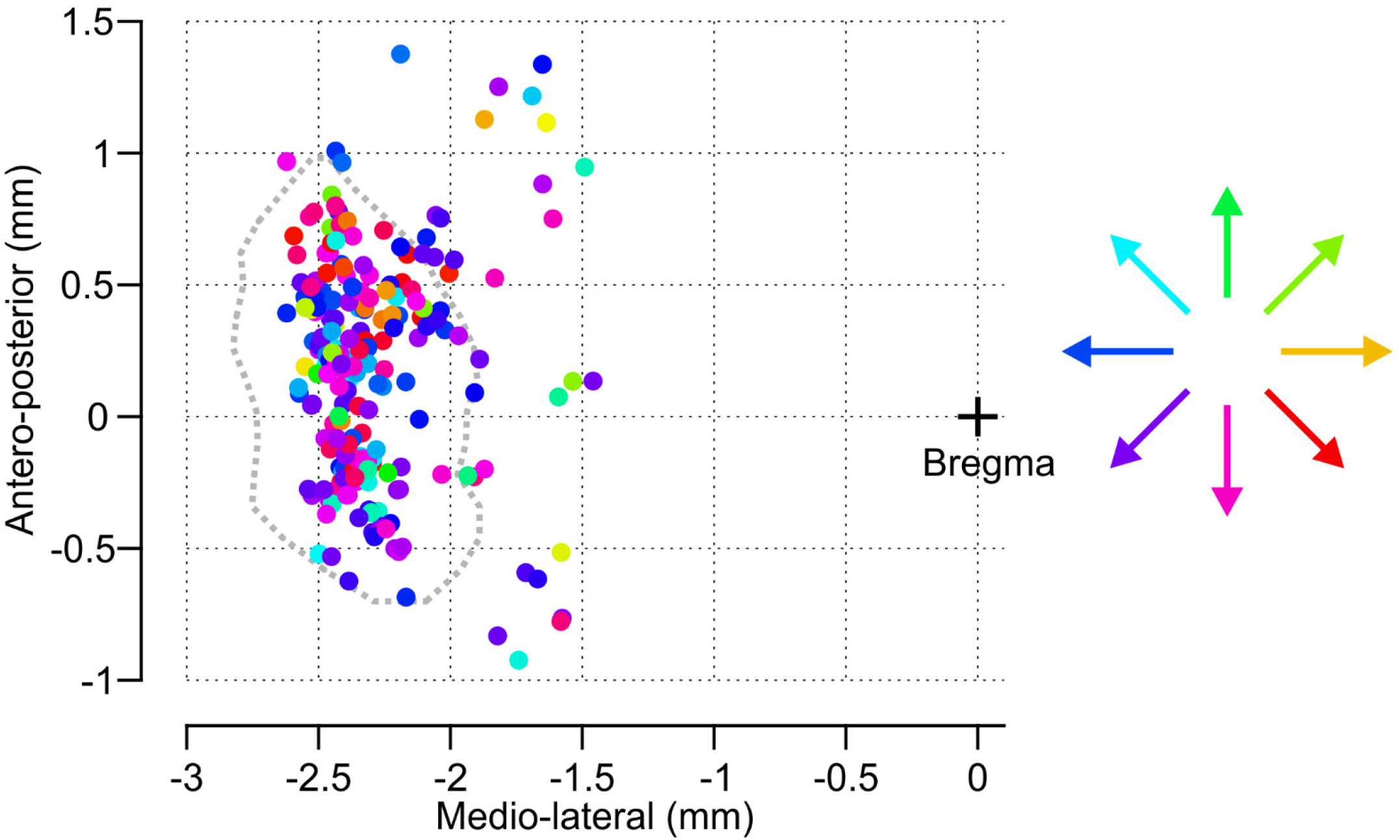
Absence of directional topography in fS1. Antero-posterior and medio-lateral coordinates relative to bregma of directionally tuned proprioceptive neurons (N=225 neurons, 17 mice). The color code corresponds to the neuron’s preferred direction. Gray dotted contour: limits of fS1 based on the mouse brain atlas^89^.

**Supplementary Movie 1. Passive forelimb displacement task.** Example trial in the passive forelimb displacement task with a robotic manipulandum.

**Supplementary Movie 2. Perceptual discrimination of proprioceptive stimuli.** Example lateral and medial displacement trials in the two-alternative forced choice (2AFC) discrimination task.

**Supplementary Movie 3. 3D tracking of joint positions.** 3D positions of 4 joints and the limb endpoint as well as their 2D projections were obtained for every manipulandum position in the tested planar workspace (red grid).

