## Supplementary Figures for "Peripersonal encoding of forelimb proprioception in the mouse somatosensory cortex"

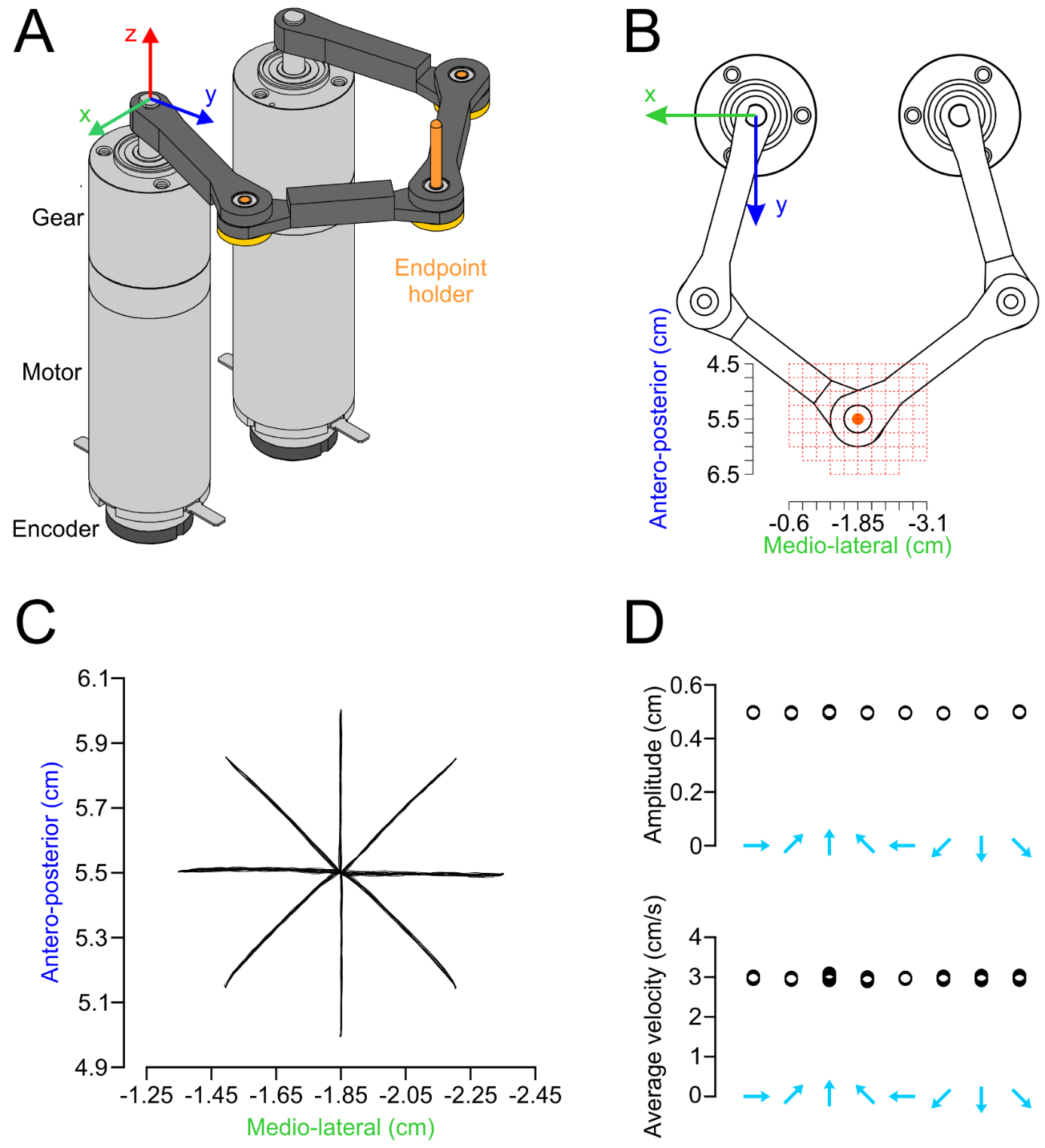

**Supplementary Figure 1. Movement kinematics of the robotic manipulandum. A**: CAD model of the robotic manipulandum showing the origin of the Cartesian coordinates. **B**: Top view of the manipulandum showing the workspace of forelimb movements (red grid). **C**: Superimposed 2D trajectories of the manipulandum’s endpoint (raw unfiltered measurements with the optical encoders and sampled at 1 kHz, see Methods) for movements in the eight tested directions in an example session (17 to 23 trials per direction, 0.5 cm amplitude and 3 cm /s velocity). **D**: Measured 2D amplitudes and average velocities of the individual movements in A were highly consistent across the tested directions.

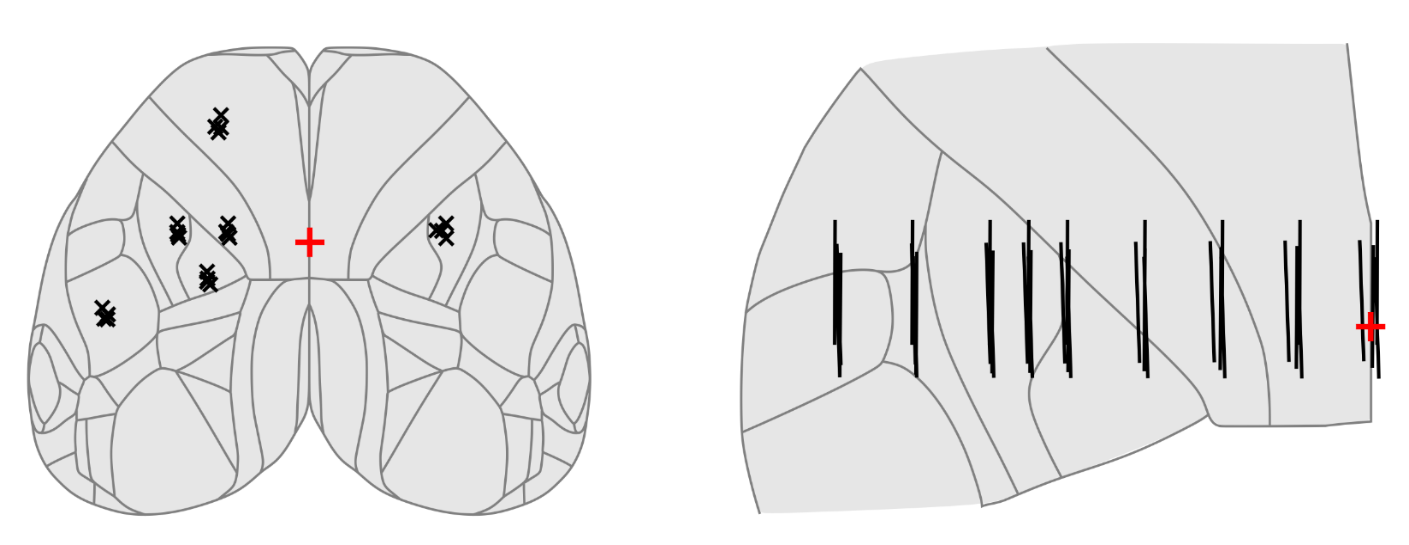

**Supplementary Figure 2. Optogenetic stimulation sites.** Stimulation coordinates of the optogenetic stimuli (Fig. 3D, F) registered to the surface projection of the Allen Mouse Brain Atlas indicate consistent targeting of the same areas across the 4 mice.

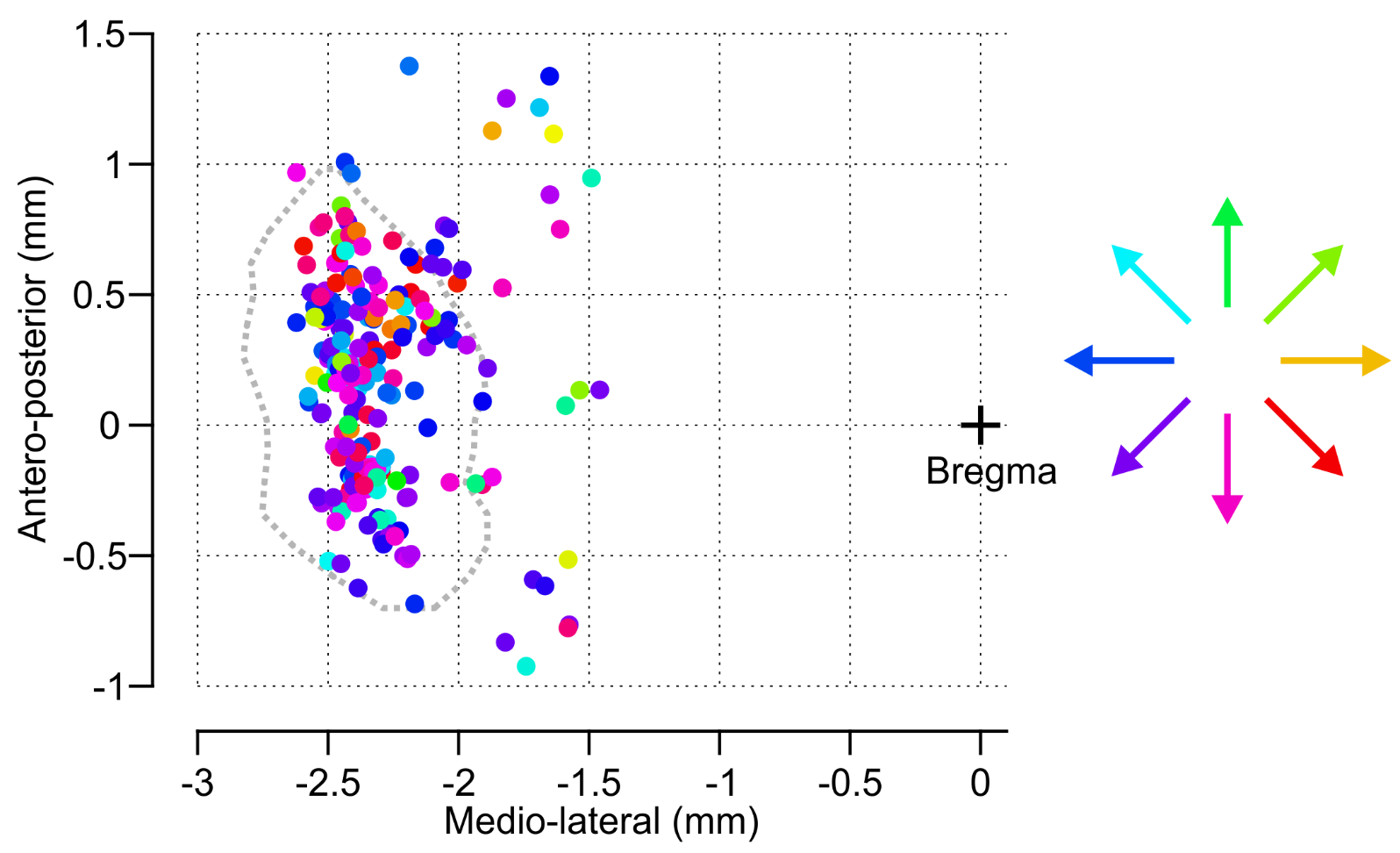

**Supplementary Figure 3. Absence of directional topography in fS1.** Antero-posterior and medio-lateral coordinates relative to bregma of directionally tuned proprioceptive neurons (N=225 neurons, 17 mice). The color code corresponds to the neuron’s preferred direction. Gray dotted contour: limits of fS1 based on the mouse brain atlas^89^.
